# Activity of zebrafish THAP9 transposase and zebrafish P element-like transposons

**DOI:** 10.1101/2024.03.22.586318

**Authors:** Nitzan Kutnowski, George E. Ghanim, Yeon Lee, Donald C. Rio

**Affiliations:** Department of Molecular and Cell Biology, University of California, Berkeley, Berkeley, CA, USA; California Institute for Quantitative Biosciences, University of California, Berkeley, Berkeley, CA, USA

**Author notes:** Correspondence: Donald Rio < >. MRC Laboratory of Molecular Biology, Francis Crick Ave. Cambridge, UK.

## Abstract

Transposable elements are mobile DNA segments that are found ubiquitously across the three domains of life. One family of transposons, called P elements, were discovered in the fruit fly *Drosophila melanogaster*. Since their discovery, P element transposase-homologous genes (called THAP-domain containing 9 or THAP9) have been discovered in other animal genomes. Here, we show that the zebrafish (*Danio rerio*) genome contains both an active THAP9 transposase (zfTHAP9) and mobile P-like transposable elements (called *Pdre*). zfTHAP9 transposase can excise one of its own elements (*Pdre*2) and *Drosophila* P elements. *Drosophila* P element transposase (DmTNP) is also able to excise the zebrafish *Pdre*2 element, even though it’s distinct from the *Drosophila* P element. However, zfTHAP9 cannot transpose *Pdre*2 or *Drosophila* P elements, indicating partial transposase activity. Characterization of the N-terminal THAP DNA binding domain of zfTHAP9 shows distinct DNA binding site preferences from DmTNP and mutation of the zfTHAP9, based on known mutations in DmTNP, generated a hyperactive protein,. These results define an active vertebrate THAP9 transposase that can act on the endogenous zebrafish *Pdre* and *Drosophila* P elements.

## INTRODUCTION

Transposable elements (TEs), or transposons, are mobile genetic elements capable of moving DNA segments from a one genomic location to another. These elements can be mobilized by either DNA or RNA intermediates, and are ubiquitous in all branches of life (Curcio and Derbyshire, 2003, Levin and Moran, 2011, Wells and Feschotte, 2020). Transposition contributes to genome evolution by causing mutations, deletions or genome rearrangements that can alter gene regulation, gene expression and gene function. Additionally, transposable element vectors are essential molecular genetic tools for manipulating genomes and for gene transfer and gene therapy (Hudecek and Ivics, 2018, Sandoval-Villegas et al., 2021). Transposable elements have also been linked to human diseases (Burns, 2020, Hickman and Dyda, 2016, Kazazian and Moran, 2017).

Many DNA-based transposable elements encode transposase proteins that catalyze the cleavage and joining reactions of transposon DNA from one genomic location to another, often using a "cut-and-paste" mechanism (Hickman and Dyda, 2015, Hickman and Dyda, 2016, Mizuuchi, 1997). Transposases are complex, multidomain proteins that catalyze transposition through a series of elaborate and consecutive assembly reactions. Structural studies on several transposases and retroviral integrases demonstrated structural and mechanistic variation within this family of proteins (Arinkin et al., 2019, Hickman and Dyda, 2015, Hickman et al., 2005). In general, transposases possess N-terminal site-specific DNA binding domains that specifically recognize a particular DNA sequence on the transposon DNA. In addition, they also contain a conserved RuvC/RNaseH catalytic domain with a so-called “DDE/D” motif catalytic, named for the three Mg^2+^-binding carboxylate catalytic amino acids in their active sites (Arinkin et al., 2019, Hickman and Dyda, 2016, Mizuuchi, 1997, Rice and Baker, 2001). A unique member of this class of enzymes, P element transposase(DmTNP), has a GTP-binding domain, uses GTP as an essential cofactor and possesses a leucine zipper dimerization domain and highly acidic carboxy-terminal domain (CTD) (Ghanim et al., 2019, Kaufman and Rio, 1992, Majumdar and Rio, 2015). DmTNP has been extensively studied (Ghanim et al., 2020, Majumdar and Rio, 2015) and a recent cryo-electron microscopy (cryo-EM) structure at 3.6 Å resolution of the DmTNP strand transfer complex (STC) revealed unique features including contact of the GTP cofactor with the transposon DNA, the arrangements of the protein domains and the unusual structures of the donor and target DNAs (Ghanim et al., 2019).

Diverse whole-genome sequencing efforts have shown that many animals, from sea urchins to humans, have a family of genes homologous to DmTNP, called THAP9. THAP9 is found in the human, zebrafish (*D. rerio*), other vertebrate and primate genomes, but THAP9 genes are partially deleted and non-functional in the mouse and rat genomes (Hagemann and Pinsker, 2001, Hammer et al., 2005, Lander et al., 2001). Previous bioinformatic analyses have demonstrated the existence of at least 50 P element-homologous sequences spread throughout the zebrafish genome called *Pdre* elements (*Pdre* [<u>P</u> homolog of *<u>D</u>anio <u>re</u>rio*]) (Hagemann and Hammer, 2006, Hammer et al., 2005). The largest one, *Pdre*2, contains 13 bp terminal inverted repeats (TIRs) together with distant 12 bp subterminal inverted repeats. The TIRs are flanked by direct 8 bp target-site duplications (TSDs), a hallmark of mobility, and notably the same size as found when *Drosophila* P elements insert in the fruitfly genome (Ghanim et al., 2020).

Here, we show that the *Drosophila* THAP9 homolog from zebrafish, zfTHAP9, encodes an active transposase protein that excise its own *Pdre* elements, as well as *Drosophila* P element transposons, despite the distinct sequences at the respective transposon termini. Additionally, DmTNP is also able to excise the zebrafish *Pdre*2 elements, regardless of the differences in the transposon termini. Based on comparisons of the known DmTNP hyperactive mutant, S129A (Beall and Rio, 1997), the recent structure of the *Drosophila* P element transposase (Ghanim et al., 2019) and multiple THAP9 sequence alignments, we identify the corresponding residue in zfTHAP9. Mutation of this residue in zebrafish THAP9 gave a hyperactive phenotype, exhibiting elevated excision activity. Taken together, our data extend previous work conducted on the vertebrate THAP9 homologs and find active P element-like *Pdre* elements in the zebrafish genome. We detected the ability of zebrafish THAP9 and DmTNP to cross mobilize their transposons. These results raise interesting questions regarding the specificity and assembly of the DNA-protein complexes required for P element family transposon mobilization.

## RESULTS

### zfTHAP9 is predicted to adopt a similar structure to the DmTNP structure

The recent cryo-EM structure of the *Drosophila* P element transposase (DmTNP) product strand transfer complex (STC) revealed unique architectural arrangements of the transposase domains and the donor and target DNAs (Ghanim et al., 2019). Based on the DmTNP structure, we carried out multiple sequence alignment (MSA) followed by structural alignment, using the T-COFFEE (Di Tommaso et al., 2011, Ghanim et al., 2019) and ITASSER (Lee et al., 2007) servers, respectively (Fig. 1A, 1B and Fig. S1).

**Figure 1.**
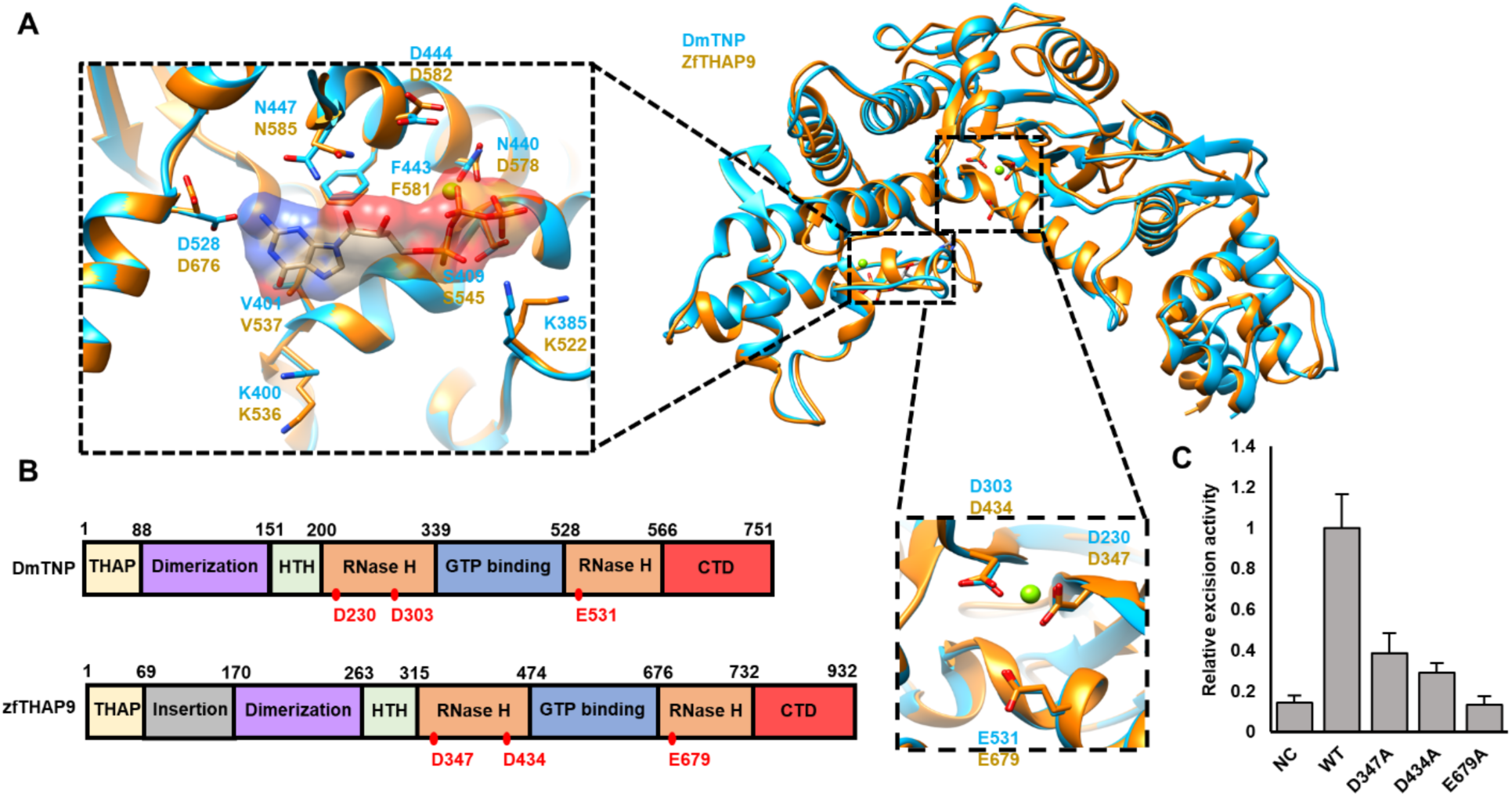
Structural alignment, domain architecture of the *Drosophila* P element transpose (DmTNP) and zebrafish THAP9 (zfTHAP9) and *Drosophila* P element excision activity of the zfTHAP9 protein. **A**. Superposition of a monomeric DmTNP cryo-EM structure (cyan) and zfTHAP9 model (gold) derived using ITASSER (Lee et al., 2007). The putative three acidic catalytic residues in zfTHAP9 and the interactions around the GTP binding site are shown. **B**. Schematic diagram of the domain architecture of DmTNP and zfTHAP9 with the boundaries indicated by amino acid residue numbers. The RNase H catalytic residues are indicated as red dots. THAP, THAP DNA-binding domain (yellow); dimerization, leucine zipper dimerization domain (purple); HTH, helix-turn-helix DNA binding domain (light green); RNase H, RNase H-like catalytic domain (orange); GTP-binding, GTP-binding insertion domain (blue); CTD, C-terminal domain (red). **C.** Results of excision assays (three replicates with standard deviation) using the *Drosophila* P elements (pISP2/Km) with the zfTHAP9 wild type and inactive mutants (D347A, D434A, E679A; NC=negative control) in *Drosophila* S2 cells (Table S2).

Overall, zfTHAP9 is predicted to adopt a similar domain organization to the DmTNP structure and other THAP9 homologs (Fig. 1A, 1B & Fig. S1). A typical THAP DNA binding domain is located at the N-terminus of the zfTHAP9 protein, like DmTNP (Fig. 1B & Fig. S1). Interestingly, an approximately 100 amino acid insertion is found in zfTHAP9 between the THAP DNA-binding and dimerization domains (Fig. 1B, Fig. S1 & Fig. S3). This unique structural feature is found only in the zfTHAP9 protein of the homologs compared (Fig. S1). Bioinformatic analysis revealed that this region is disordered and was unable to predict a well-folded structure domain for this insertion (Fig. S3). We also generated models of the zfTHAP9 dimer using Alphafold2 (Liu et al., 2023, Mirdita et al., 2022, Mirdita et al., 2019). Analysis with Alphafold2 also suggested this region is disordered. Given the distinctive organization of the zebrafish *Pdre*2 elements (Hammer et al., 2005), this unique insertion may act as a structural linker that helps the protein to engage with the remote internal and terminal inverted repeats in order to promote DNA-protein assembly for catalysis of transposition. Most interestingly, there is a critical difference between the Alphafold2 model of zfTHAP and the cryo-EM structure of the *Drosophila* P element transposase (Ghanim et al., 2019) (Fig. 2 & Fig. S2). In the zfTHAP9 model, two alpha helices are predicted to occupy the same position that the target DNA does in the *Drosophila* transposase strand transfer structure (Fig. S2C-D) that occupy the same position as the target DNA does in the *Drosophila* transposase structure (Fig. S2C-D). This predicts that zfTHAP9 may not be able to carry out forward transposition or that strand transfer into a target DNA by zfTHAP9 would require a conformational change of these helixes. This observation may explain why the zfTHAP9 protein exhibits *Pdre* element excision but is uncapable of forward transposition (see below).

**Figure. 2:**
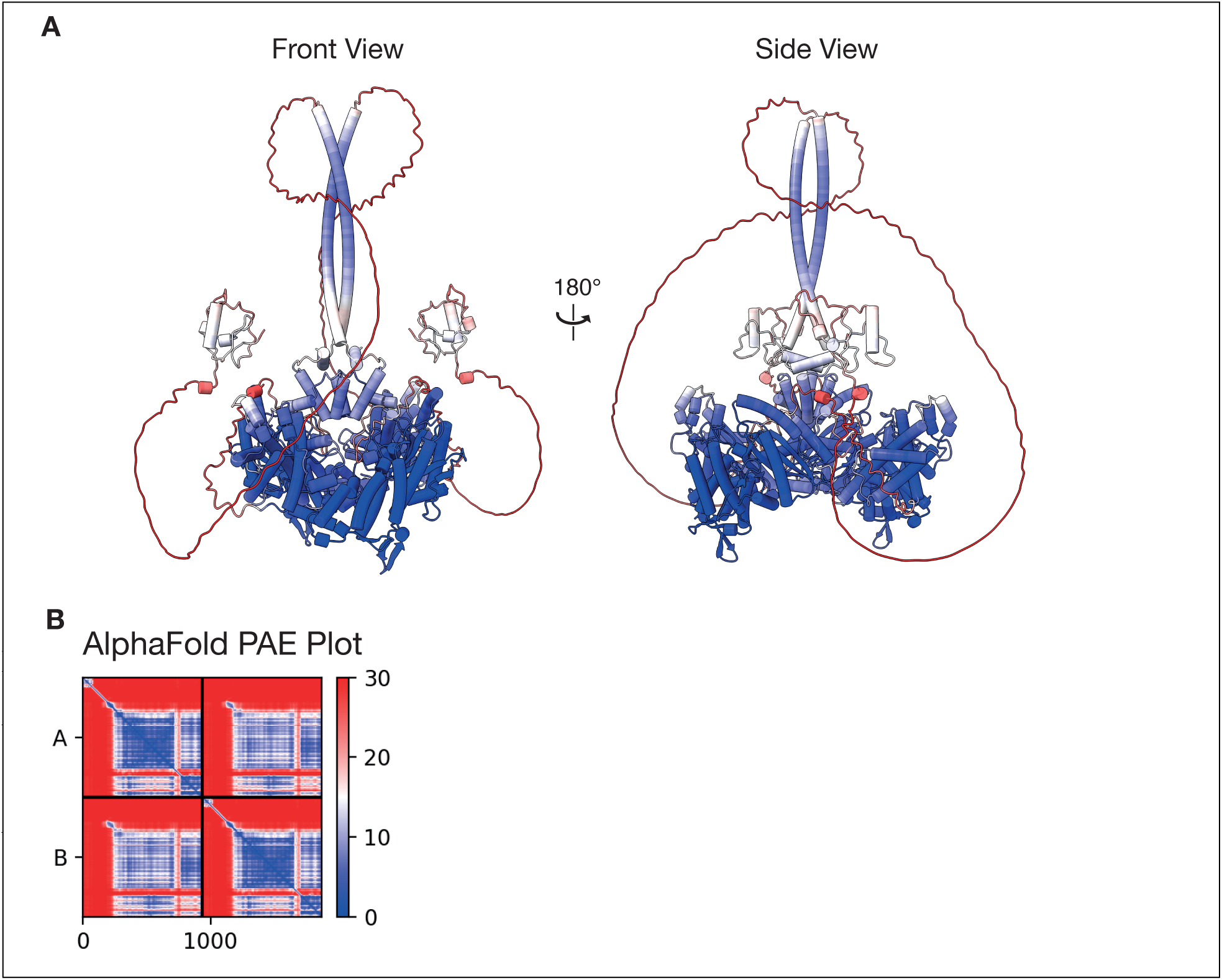
AlphaFold Prediction and PAE plot of zfTHAP9 dimer. **A.** Front and side views of an AlphaFold prediction of the zfTHAP9 dimer. Residues are coloured according to PAE confidence values. **B.** AlphaFold PAE plot of the zfTHAP9 AlphaFold prediction.

Based on the DmTNP structure, we generated structural models for zfTHAP9 using the ITASSER (a.a. 239-719) (Lee et al., 2007). Structural alignment of the model with the DmTNP structure is consistent with the MSA result (Fig. 1 & Fig. S1). The THAP DNA binding domain and most of the leucine zipper dimerization region of DmTNP were too flexible and therefore were not resolved in the cryo-EM structure. Thus, our zfTHAP9 model begins with the N-terminal DNA binding helix-turn-helix domain followed by a split catalytic RNase H domain (RNaseH) that is interrupted by a GTP-binding insertion domain (GBD) and a C-terminal domain (CTD).

The RNaseH catalytic domain contains a DDE motif that, together with Mg^2+^, carries out the transposition reaction (Ghanim et al., 2019). The catalytic residues deduced from the zfTHAP9 model (D347, D434 and E679) aligned well with those of DmTNP (Fig. 1A & Table S1). The DmTNP RNaseH catalytic domain is disrupted by the insertion of a GTP binding domain, like insertions found in several other transposases (Ghanim et al., 2019, Goryshin and Reznikoff, 1998) (Fig. 1, Fig. S1 & Table S2). DmTNP requires GTP as a cofactor, without hydrolysis, for the assembly of the active protein-DNA synaptic complex (Mul and Rio, 1997, Tang et al., 2005). The residues that interact with the GTP in DmTNP are conserved in zfTHAP9 suggesting a similar interaction pattern and function for GTP in zfTHAP9 (Table S2). Furthermore, DmTNP K400 in that interacts with the transferred strand DNA bases is conserved in zfTHAP9 (K536), reinforcing the idea that zfTHAP9 employs a similar DNA transposition mechanism (Fig. 1A, Fig. S1 & Table S2).

To validate the computational predictions of the putative catalytic residues in zfTHAP9, each residue (D347, D434 and E679) was substituted to alanine and tested for activity using excision assays in *Drosophila* cells. Indeed, alanine substitution of any one of these acidic residues dramatically reduced zfTHAP9 excision activity *in vivo*, confirming their essential role for transposase catalytic activity (Fig. 1C & Table S3). Similar results were obtained with substitution of the corresponding DmTNP catalytic residues to alanine (Ghanim et al., 2019).

### zfTHAP9 can excise the zebrafish *Pdre*2 elements and *Drosophila* P elements

Previous bioinformatic work demonstrated the existence of at least 50 P-homologous sequences spread throughout the zebrafish genome called *Pdre* elements (*Pdre* [<u>P</u> homolog of *<u>D</u>anio <u>re</u>rio*]). In addition to the full-length *Pdre*2 THAP9 transposase-like gene, there are also multiple internally-deleted *Pdre* elements elsewhere in the zebrafish genome. *Pdre*2 is located on linkage group 2 (accession number BX511023), contains 13 bp terminal inverted repeats (5’-CATACCTGTCAAC-3’and 5’-GTTGACAGGTATG-3’) together with 12 bp subterminal inverted repeats (5’-TGTTTAAACCAA-3’and 5’-TTGGTTTAAACA-3’) and is flanked by direct 8 bp target-site duplications (TSD) (5’-AGGTGAAT-3’), a hallmark of mobility (Hagemann and Pinsker, 2001, Hammer et al., 2005). Therefore, we cloned the zfTHAP9 ORF into expression vectors suitable for expression in either human HEK293 or *Drosophila* S2 cells. To investigate the excision activity of zfTHAP9 transposase, we cloned the full-length zebrafish *Pdre*2 elements into a reporter plasmid, termed pISP/*Pdre*2, that was similar to the pISP-2/Km reporter in previous studies on Drosophila P element excision (Rio et al., 1986).

zfTHAP9 transposase expresses in both human HEK293 and *Drosophila* S2 cells, as indicated by immunoblotting (Fig. S4). We then tested zfTHAP9 excision activity using *in vivo* excision assays in either *Drosophila* S2 and human cells (Fig. 3A). The excision assay results showed that zfTHAP9 excises its own *Pdre*2 element and also the *Drosophila* P element transposon in both cell types (Fig. 3B-F & Fig. S5-S7), yet with higher efficiency in *Drosophila* cells. These data suggested that zfTHAP9 acts as an active transposase protein that cleaves both elements even though the repeat organization between the elements is different.

**Figure 3.**
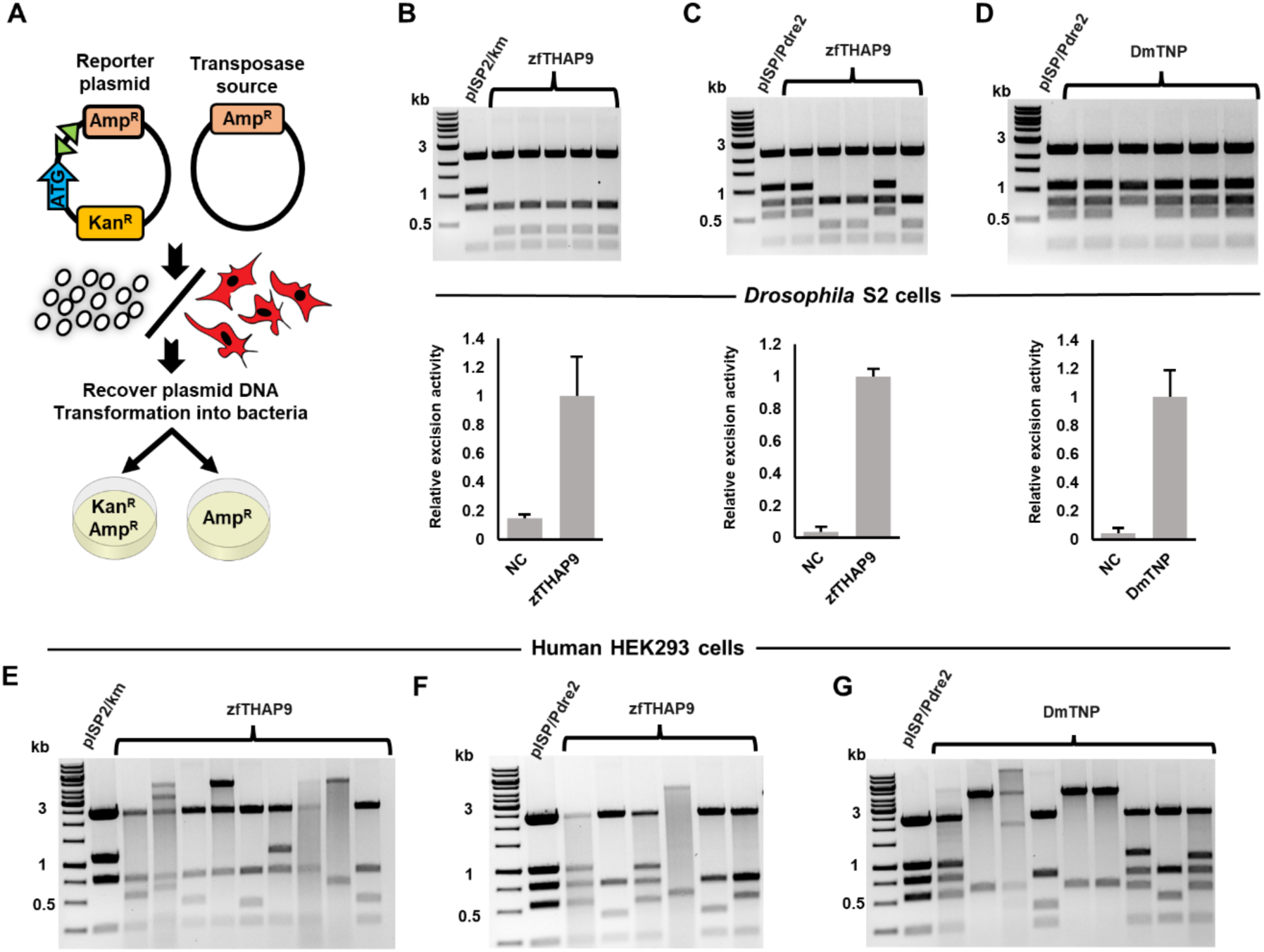
*Drosophila* P element and zebrafish *Pdre*2 element excision assays with zfTHAP9 and DmTNP in *Drosophila* S2 and human HEK293 cells. **A.** Diagram of the excision assay. **B-D.** Upper panel: Images of ethidium bromide-stained 1% agarose gels showing the *Pvu*II cleavage patterns of the pISP2/Km or pISP/*Pdre*2-derived plasmid DNAs that were recovered from single kanamycin and ampicillin-resistant bacterial colonies obtained after transfection of *Drosophila* S2 cells and electroporation of *E. coli*. Lower panel: graphs of the relative excision activity (three replicates with standard deviation, NC=negative control). **B.** zfTHAP9 + pISP2/Km; **C.** zfTHAP9 + pISP/*Pdre*2 and **D.** DmTNP + pISP/*Pdre*2 (Table S2). **E-F.** Images of ethidium bromide-stained 1% agarose gels showing the *Pvu*II cleavage patterns of the pISP2/Km or pISP/*Pdre*2-derived plasmid DNAs that were recovered from single kanamycin and ampicillin-resistant bacterial colonies obtained after transfection of HEK293 cells and electroporation of *E. coli*. **E.** zfTHAP9 + pISP2/Km; **F.** zfTHAP9 + pISP/*Pdre*2 and **G.** DmTNP + pISP/*Pdre*2.

While the excision events of *Drosophila* P elements showed small flanking deletions with DmTNP, larger deletions of donor DNA flanking the two transposon ends were observed after excision by zfTHAP9 in *Drosophila* S2 cells (Fig. S6). This may be related to the ability of the zfTHAP9 to quickly release the donor DNA ends after cleavage. If the protein is loosely bound to the donor DNA, endonucleases could digest the DNA before repair by the DNA repair machinery in the cells which would lead to larger deletions compared to DmTNP.

Excision assays indicate that both DmTNP and zfTHAP9 can excise the zebrafish *Pdre*2 elements in both cell types (Fig. 3, Fig. S5-S7). The *PvuII* digestion pattern of the excised *Pdre*2 reporter plasmid showed an “unexcised” or “uncut” pattern in some samples. Sequencing of these samples showed cutting or small deletions at one transposon termini. These cleavage events are small and is hard to detect on an agarose gel, yet can restore an open reading frame for kanamycin in the reporter plasmid (Fig. S7). This suggests that the two *Pdre*2 element ends are cleaved asynchronously. The same cleavage pattern has been observed in *Drosophila* P elements (Tang et al., 2007). No such DNA repair products were observed in the negative control samples (data not shown).

Notably, all excision assays in HEK293 cells yielded a large portion of samples exhibiting aberrant DNA rearrangements after DNA excision (Fig. 3, Fig. S5-S7). A possible explanation could be related to how the proteins function in the two cell types. Differences in DNA repair or recombination mechanisms or cellular components that may results in differential interactions in human cells that could give rise to the aberrant products. Furthermore, the total number of ampicillin-kanamycin resistant colonies was dramatically lower in human HEK293 cells compared to *Drosophila* S2 cells, suggesting that the activity of the two proteins is lower in human cells. Taken together, the excision assay results showed that both transposases can cleave the *Drosophila* P elements as well as zebrafish *Pdre*2 elements in both cell types, likely via engagement of the THAP DNA-binding domain and other DNA binding regions of the proteins with the transposon DNA.

### Assays to detect transposition of *Pdre*2 elements by zfTHAP9

To understand if zfTHAP9 can integrate transposon DNA, we performed plasmid-to-plasmid transposition assays in *Drosophila* S2 and human HEK293 cells. As a positive control, we tested integration of the *Drosophila* P elements by DmTNP. In this experiment we co-transfected the cells with a transposase source, a donor plasmid, which contains a tetracycline resistance gene between the *Drosophila* P element or the zebrafish *Pdre*2 element ends and a target plasmid (Fig. 4A). A GFP expression plasmid containing an ampicillin resistance gene was used as a target plasmid. One day after transfection, the plasmids were extracted from the cells which then yielded tetracycline and ampicillin-resistant bacterial colonies in and followed by sequencing, which indicated integration of the *Drosophila* P elements by DmTNP into the GFP plasmid. All the integration events occurred in the promoter or in the GFP gene (Fig. 4B-D). These results are consistent with previous work showing that P elements insertions often occur at regions around gene promoters and at regions overlapping origins of DNA replication or open chromatin (Remus et al., 2004, Spradling et al., 2011).

**Figure 4.**
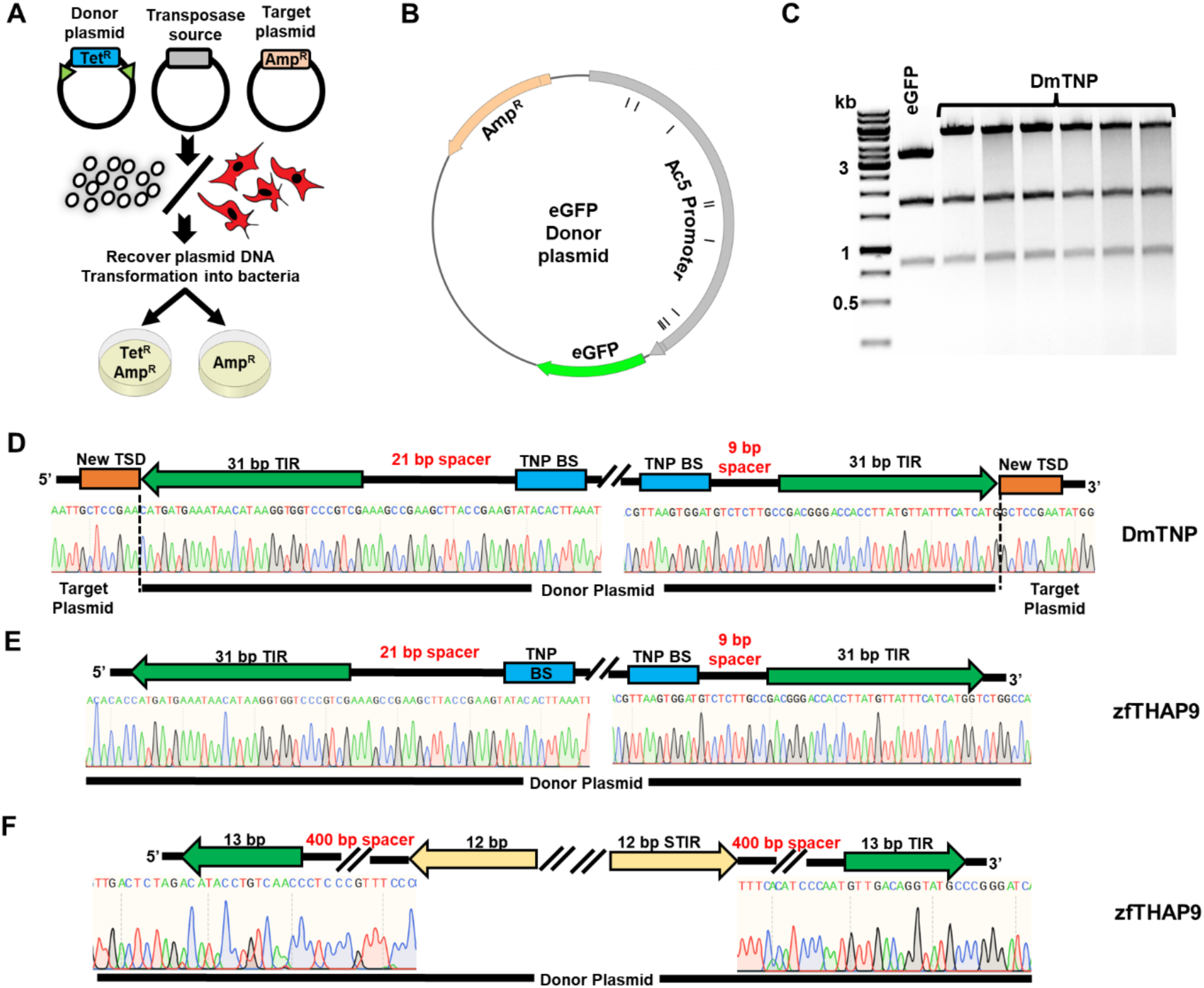
Plasmid-to-plasmid transposition assay. **A.** Diagram of plasmid-to-plasmid transposition assays. **B.** The map of target eGFP plasmid is shown. The black lines indicate locations of the P element new target site duplications (TSDs) after integration in S2 cells. **C.** *BsaI* cleavage pattern for plasmid DNAs that were recovered from single tetracycline + ampicillin-resistant bacterial colonies obtained after transfection of *Drosophila* S2 cells with the DmTNP expression plasmid. The increase of 1.7 kb to the upper band compared to the donor plasmid is indicative of forward transposition events. **D-F.** Raw sequencing data of plasmid DNA after a new integration in *Drosophila* S2 cells. The boundaries of the donor plasmid are shown with the organization of the *Drosophila* P elements. The new TSDs that are part of the transposed target plasmid are colored in orange. **D.** DmTNP-S129A + pISP2/Km **E.** zfTHAP9 + pISP2/Km; **F.** zfTHAP9 + pISP/*Pdre*2.

While DmTNP can integrate its own elements in *Drosophila* S2 cells, plasmid-to-plasmid assays with zfTHAP9 yielded no transposition events. Instead, only nonspecific DNA rearrangement events were recovered in which no TSDs were detected (Fig. 4E). In addition, no integration events were observed of the zebrafish *Pdre*2 element with either protein (Fig. 4F). These results indicate that while zfTHAP9 and DmTNP can excise the *Pdre*2 transposon DNA, zebrafish THAP9 cannot carryout the forward transposition reaction to integrate the excised transposon DNA. The absence of forward transposition might be due to lack of a zebrafish protein cofactor that stimulates zfTHAP9 transposition activity, which would be absent in our *Drosophila* S2 or human HEK293 cellular assays. Alternatively, the zfTHAP9 protein may be unable to bind target DNA as predicted by the clash between helixes of the zfTHAP9 AlphaFold model and target DNA in the Drosophila P element transposase strand transfer structure.

The observed nonspecific DNA rearrangement events may arise from the broken DNA ends generated by zfTHAP9 undergoing nonspecific nuclease activity and those ends can be joined to the ampicillin resistant plasmid to give rise to both tetracycline and ampicillin resistance plasmids, unlike the bona fied transposition products observed with DmTNP. We call this nonspecific activity “random integration”. Plasmid-to-plasmid experiments performed in human cells showed no transposition events, even with the DmTNP and *Drosophila* P elements, suggesting that while transposon excision can occur, forward transposition does not occur in human cells using this assay.

We next tested the ability of the proteins to mobilize *Drosophila* P elements or zebrafish *Pdre*2 elements, genetically marked with a puromycin selection marker, from a plasmid into the human genome. For that we used integration assays followed by splinkerette PCR (spPCR) (Potter and Luo, 2010). Integration assay results revealed significantly higher number of puromycin-resistant colonies with each one of the proteins over the negative control (Fig. S8). However, spPCR analysis of individual isolated colonies indicated that only “random integration” events occurred. These results are consistent with the plasmid-to-plasmid assay results and suggest that in human cells both proteins failed to integrate either transposons and that the cleaved transposon DNA integrates by a nonspecific integration mechanism.

Next, we conducted plasmid rescue experiments, a well-established method for recovering transposon integrations (Beckermann et al., 2021, Huang et al., 2009). This method involves no PCR amplification and therefore, is not subject to potential PCR artifacts or ligation of PCR-amplified products (Fig. 5A). The plasmid rescue results agreed with the plasmid-to-plasmid assay results, indicating DmTNP can excise and integrate its own elements into the genome in *Drosophila* S2 cells (Fig. 5B-C). However, no integration events were detected with zfTHAP9 with either *Pdre*2 or *Drosophila* P elements in human HEK293 or *Drosophila* S2 cells. We observed clear transposition activity for DmTNP only in *Drosophila* S2 cells. The colonies arising from this assay may be due to excision of the transposon or cleavage at one end and random integration that is dependent on DmTNP or zfTHAP9 activity, since only a few colonies are obtained without including a transposase expression plasmid (Fig. S8).

**Figure 5.**
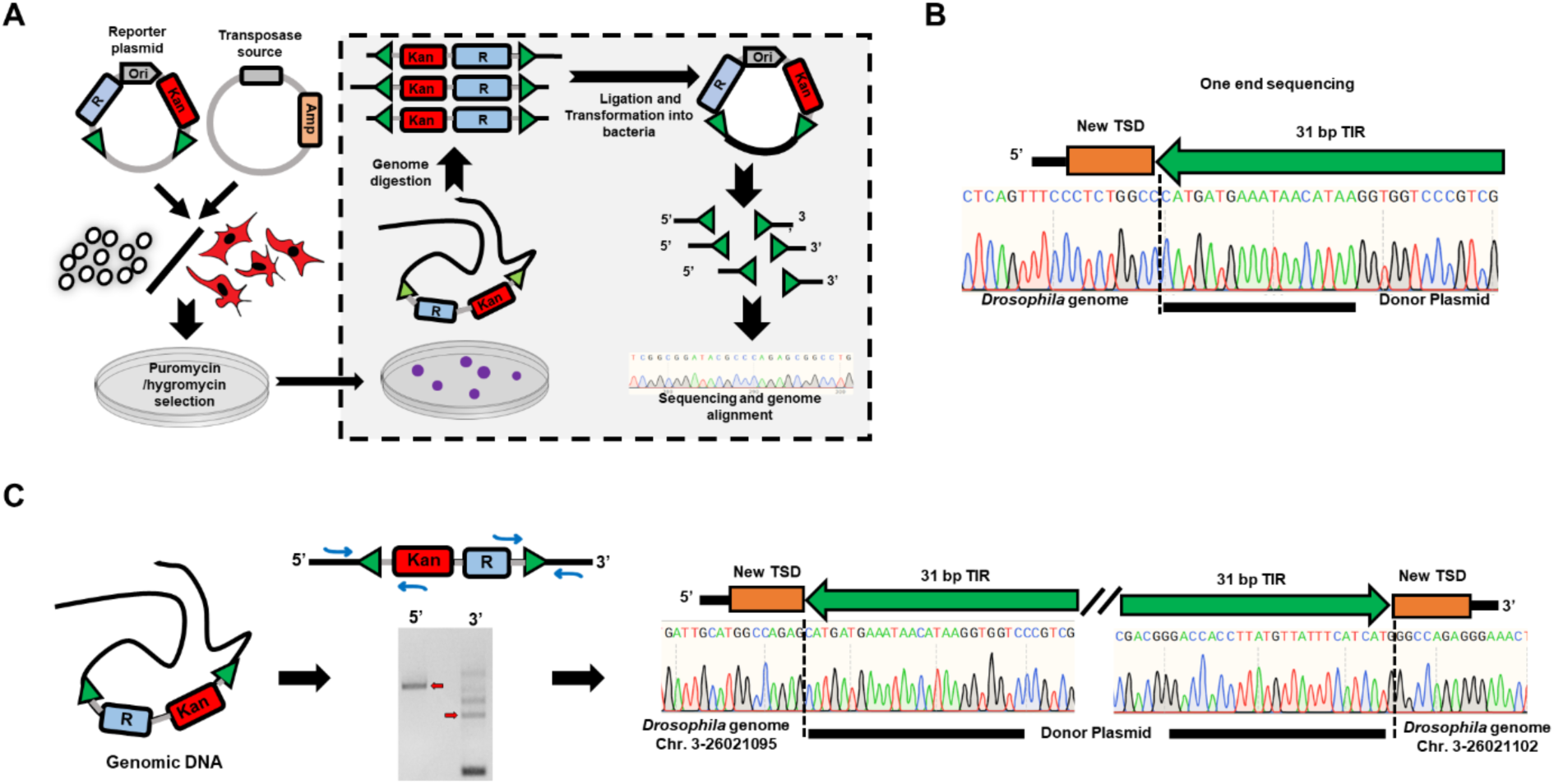
Rescue of integrated P elements plasmid DNA from *Drosophila* cells. **A.** Diagram of plasmid-to-genome integration assays. **B.** Raw sequencing data of P elements-containing plasmid DNA recovered after genomic integration into the *Drosophila* genome. The terminal inverted repeats (TIRs) of the donor P elements are shown with the new TSDs that are part of *Drosophila* genome the colored in orange. **C.** Left, PCR using *Drosophila* genomic as a template DNA to verify transposition. The primer design was based on the genomic location obtained from sequencing results of one transposon end. Right, PCR products were cut from the gel, cloned and sequences were determined by Sanger sequencing. The TIRs of the donor plasmid are shown with the new TSDs that are part of the *Drosophila* genome colored in orange.

### Identification and analysis of hyperactive and inactive mutants

Previous work on DmTNP found one specific mutation, S129A, that resulted in an elevated level of transposase activity in *in vivo* recombination assays, including P element mediated germline transformation (Beall et al., 2002). *In vitro* assays for P element transposase activity indicated that the S129A mutant exhibits elevated donor DNA cleavage activity when compared to the wild-type protein, whereas the strand-transfer activity is similar to that of wild type (Beall et al., 2002).

Inspection of the DmTNP structure suggested that this residue S129 is found in the leucine zipper dimerization domain and was the first amino acid traced in the cryo-EM map (Fig. 6A-B). Based on this hyperactive DmTNP mutant, we performed MSA analysis with the zebrafish and other THAP9 homologs. We found an equivalent position, S239 in the zfTHAP9 sequence (Fig. 6A). Superposition of zfTHAP9 model with the DmTNP structure suggested a similar location for this residue (Fig. 6B-D). To examine this residue affects activity, we substituted this equivalent position to alanine in zfTHAP9 and tested activity using excision and plasmid-to-plasmid assays. Since many aberrant DNA rearrangements were observed in HEK293 cells, we tested the activity of the mutants only in *Drosophila* S2 cells.

**Figure 6.**
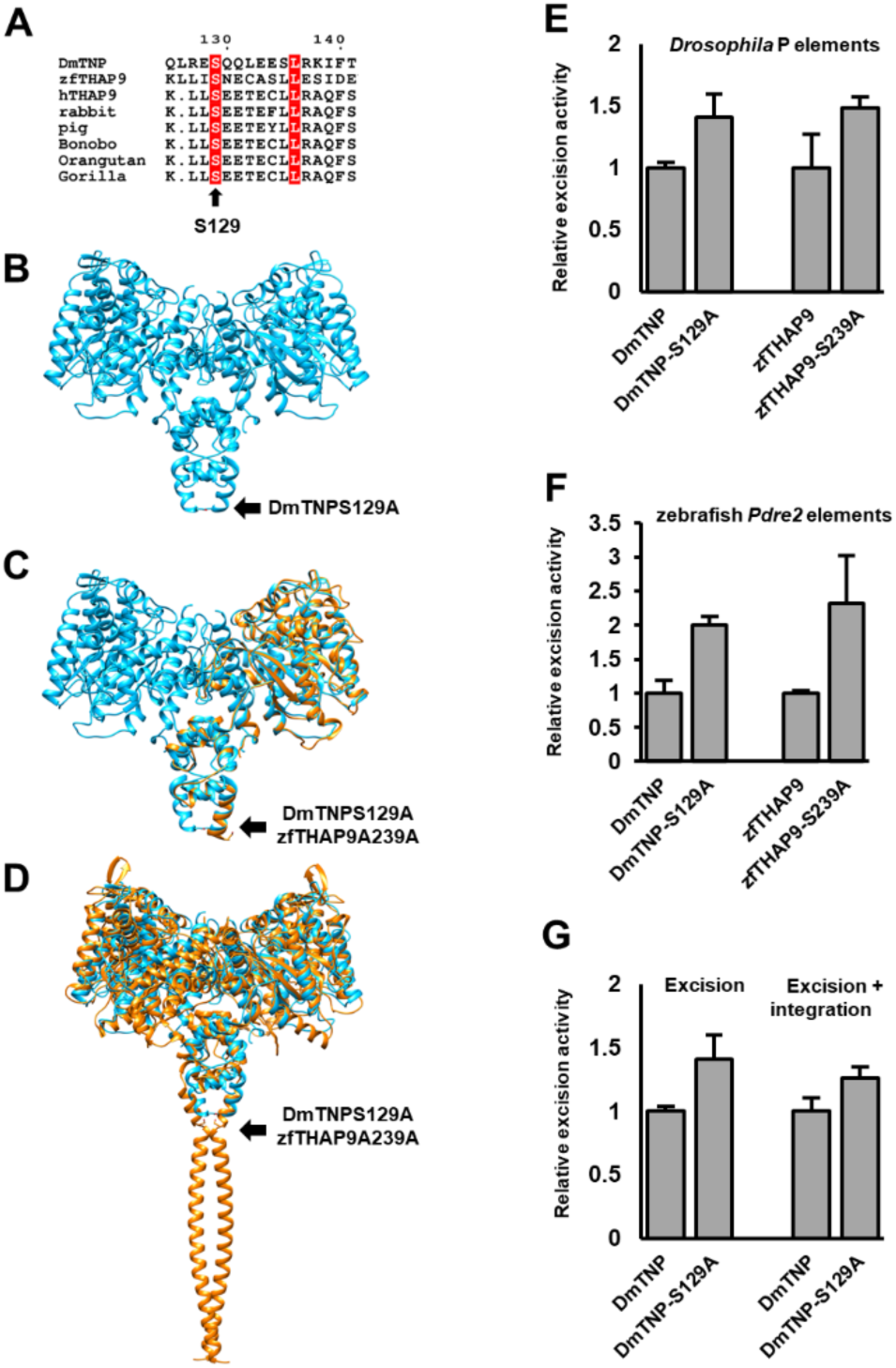
Mutations in DmTNP and zfTHAP9 that elevate excision activity. **A.** Multiple sequence alignment (MSA) of *Drosophila* P element transposase (DmTNP) with related THAP9 proteins. THAP9 proteins: *Drosophila* P element transposase (DmTNP), zebrafish THAP9 (zfTHAP9), human THAP9 (hTHAP9) with homologous proteins from rabbit, pig, bonobo, orangutan and gorilla. Amino acids in the red boxes are identical. The sequences were aligned using Tcoffee (Di Tommaso et al., 2011) and displayed as determined by ESPript3.0 (Robert and Gouet, 2014). Position S129 of DmTNP is marked. **B.** DmTNP structure showing the location of S129. **C.** Superposition of DmTNP cryo-EM structure (cyan) and zfTHAP9 model (gold). Positions S129 of DmTNP and S239 of zfTHAP9 are marked by black arrows. **D.** Superposition of the DmTNP cryo-EM structure (cyan) and zfTHAP9 model (gold), including the dimerization domain. Positions S129 of DmTNP and S239 of zfTHAP9 are marked. All zfTHAP9 models derived from Alphafold2 (Jumper et al., 2021). **E.** Excision assay results of the *Drosophila* P element with the wild type and mutant proteins, DmTNP S129A and zfTHAP9 S239A in *Drosophila* S2 cells. **F.** Excision assay results of the zebrafish *Pdre*2 element with the wild type and mutant proteins in *Drosophila* S2 cells. **G.** Excision and plasmid-to-plasmid integration assays results for the *Drosophila* P element with the DmTNP wild type and hyperactive mutant in *Drosophila* S2 cells.

The wild-type and mutant zfTHAP proteins expressed to similar levels (Fig. S4). In agreement with previous work (Beall et al., 2002), DmTNP S129A showed elevated excision and integration activity of the *Drosophila* P elements compared to wildtype DmTNP (Fig. 5E-G). DmTNP S129A can also efficiently excise zebrafish *Pdre*2 elements (Fig. 5F). Like DmTNP S129A, the zfTHAP9 S239A mutant showed elevated excision activity and acts as a hyperactive mutant (Fig. 6E-F). However, no integration events were detected with the zfTHAP9 S239A as observed with wildtype zfTHAP9. These results demonstrated that a single point mutation in the leucine zipper dimerization domain, which links the DNA binding THAP domain to the catalytic domain, can enhance protein transposition activity.

### DNA binding site preferences for the zfTHAP9 THAP domain

Transposases often possess N-terminal site-specific DNA binding domains that specifically recognize particular DNA sequences on the transposon DNA (Arinkin et al., 2019, Hickman and Dyda, 2015, Hickman and Dyda, 2016, Mizuuchi, 1997, Rice and Baker, 2001). The *Drosophila* P elements contain an internal 10 bp high-affinity DNA binding site at each end, that are initially recognized by the THAP DNA-binding domain (Tang et al., 2005). However, no similar sequences to the P element DmTHAP domain binding sites are found in the zebrafish *Pdre*2 sequence, yet excision events were observed by DmTNP in both cell types (Fig. S5-7).

To find high affinity recognition motifs of zfTHAP9, we expressed and purified the zfTHAP9 THAP DNA-binding domain (1-69 a.a.) to carry out *in vitro* selection (SELEX) experiments with random oligonucleotide pools. After ten rounds of enrichment, we found an over-represented sequences that converged to a defined motif (Fig. 7). This motif was not similar to the *Drosophila* P element THAP binding sites. Interestingly, no similar motifs were found in the zebrafish *Pdre*2 element, which raises questions of how the zfTHAP domain engages with *Pdre*2 and P element DNA. It is possible that a series of lower affinity DNA binding motifs may allow zfTHAP9 to assemble on and excise the *Pdre*2 and P element transposon DNAs.

**Figure 7.**
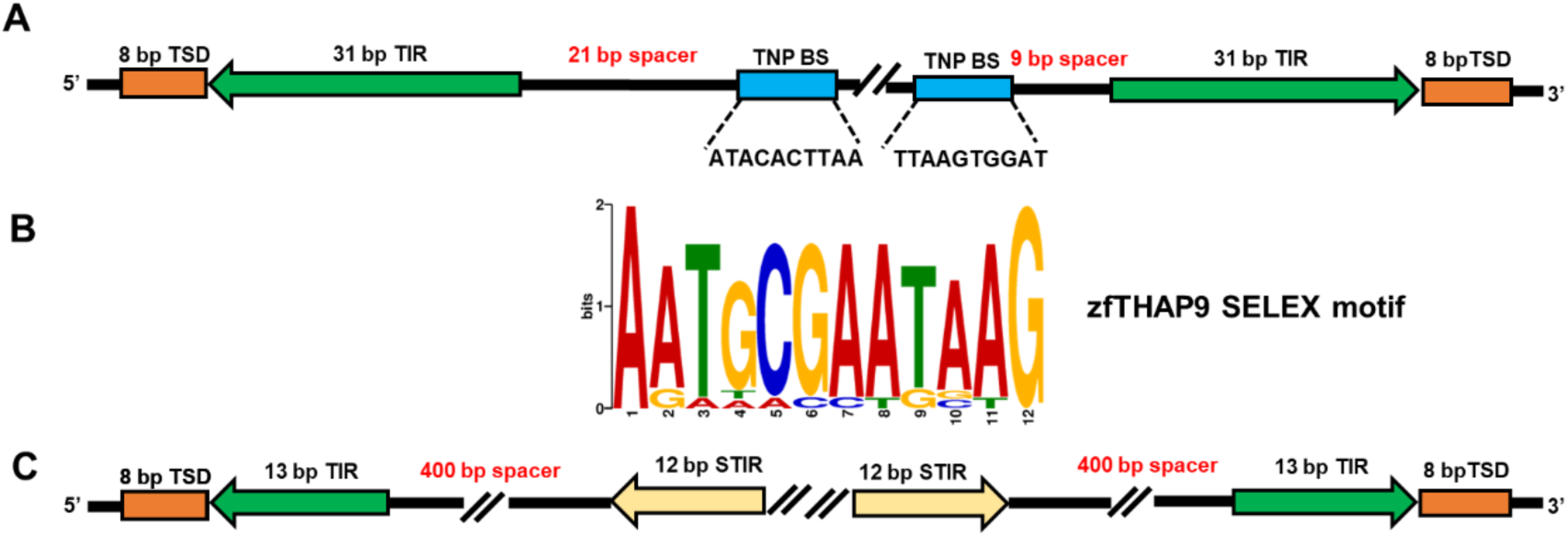
Drosophila P element and zebrafish transposon organization and the zfTHAP9 DNA binding SELEX results. **A.** Diagram of the *Drosophila* P elements including the sequences of the known DmTNP binding sites. **B.** Zebrafish THAP domain DNA binding SELEX motif generated using MEME (Bailey et al., 2009). **C.** Diagram of the zebrafish *Pdre*2 element.

## Discussion

P elements are one of the most extensively studied eukaryotic DNA transposons (Ghanim et al., 2020). Biochemical and structural studies have given a wealth of insights into the cellular regulation and mechanism of transposition and the details of the transposase nucleoprotein complex (Ghanim et al., 2020). Like other autonomous DNA-based transposons, P elements encode a transposase, an enzyme that is responsible for DNA transposition. Transposition occurs via a defined series of consecutive steps: transposase-transposon DNA binding, pairing of the transposon ends (synaptic complex formation), donor DNA cleavage, target DNA capture, strand transfer (integration) and disassembly and DNA repair. In general, transposition can be divided into two reactions: DNA cleavage and DNA integration (Ghanim et al., 2020, Hickman and Dyda, 2015). DmTNP exhibits unique transposase features, such as the use of GTP as a required cofactor for the DNA pairing, cleavage and strand transfer stages of transposition (Ghanim et al., 2020, Kaufman and Rio, 1992). In addition, the staggered cleavage at the P element transposon ends is atypical, resulting in a 17 nt 3′-single-stranded transposon DNA extension (Beall and Rio, 1997, Ghanim et al., 2019, Kaufman and Rio, 1992).

In this work, we showed that the *Drosophila* P element transposase homolog from zebrafish, zfTHAP9, encodes an active protein that is able to excise its own elements, *Pdre*2 and *Drosophila* P elements. We also extended our knowledge regarding the activity of DmTNP bydistinguish the excision and integration steps during transposition *in vivo*. DmTNP can excise the zebrafish *Pdre*2 elements, although the TIRs and IIRs are organized differently from the *Drosophila* P elements (Fig. 7). Based on earlier studies (Kaufman et al., 1989), this process may involve recognition and binding of the THAP DNA binding domain to one transposon end to initiate the capture and pairing of the second end, in a GTP-dependent manner, to form the paired-end complex (Tang et al., 2005). When the THAP domains of DmTNP engage with the internal 10 bp transposase binding sites and internal IR binding sites, then DmTNP acts to pair the two different P element ends in a manner thought to be similar to the 12–23 rule imposed by the RAG1–RAG2 V(D)J recombinase (Ghanim et al., 2019, Kim et al., 2018, Lapkouski et al., 2015, Ru et al., 2015). Hence it is interesting how the DmTNP THAP domain interacts with the zebrafish *Pdre*2 elements to initiate the excision reaction. Moreover, SELEX results suggest a different high-affinity binding motif for the zfTHAP domain. It is possible that DNA shape or flexibility, rather than strict DNA sequence impact the interaction and assembly of zfTHAP9 to initiate transposition, like the bacterial transposon Tn7 when TnsB interacts with DNA (Kaczmarska et al., 2022). There is evidence that DmTNP can, at elevated concentrations, coat a P element end DNA in footprinting assays (Kaufman et al., 1989).

Previous studies with full-length P element ends indicated that a transposase tetramer acts at the early stages of transposition in forming synaptic paired end (PEC) and cleaved donor (CDC) complexes (Tang et al., 2007, Tang et al., 2005). However, a cryo-EM structure showed that transposase is dimeric in the strand transfer complex (STC)(Ghanim et al., 2019). Therefore, it is possible that as the transposition reaction proceeds, different oligomeric states are needed to accommodate each stage. For example, a tetramer (or a dimer of dimers) initially assembles to pair the natural P element ends and activate the protein for DNA cleavage. Once the P element ends are excised and the target DNA is captured, only two catalytic subunits would be required to form the strand transfer complex.

In DmTNP, S129 is located at the end of the leucine zipper-dimerization domain. Substitution of S129 to alanine enhances DNA cleavage and integration (Fig. 6). These results emphasize the idea that large structural transitions and rearrangements must occur at the P element transposon ends to carry out transposition. Surprisingly, substitution of the equivalent position to S129 in the zfTHAP9 sequence, S239 to alanine also yielded a hyperactive protein. Multiple sequence alignments (MSA) suggested that this position is fully conserved among THAP9 genes (Fig. 6A). It is possible that the S129A mutant may alter interactions important for maintaining the oligomeric form of the protein, which might affect the ability of the active sites to engage with the transposon ends, leading to a more active protein.

In this work we showed that DmTNP can excise the *Drosophila* P elements, but no integration events were detected using three different methods in human cells, unlike in *Drosophila* S2 cells. Previous studies showed that purified DmTNP can catalyze transposition *in vitro* (Ghanim et al., 2019, Kaufman and Rio, 1992). These results together raise the possibility that in human cells, other proteins or molecules can interact after P elements excision and dramatically reduce the transposition to an undetectable level. Another explanation may be that P elements get excised in a non-concerted reaction where one end gets cut first and then the other (Ghanim et al., 2019, Kaufman and Rio, 1992, Tang et al., 2005), and this could lead to random integration into the human genome.

The structural analysis also illuminates and may explain why zfTHAP9 cannot complete transposition. There are a pair of interacting alpha helices near the C-terminus of zfTHAP9 (R920-F929), where each α helix belongs to a different subunit in the dimeric structure (Fig. S2A-C). Based on DmTNP structure, this structural element may contribute to the dimerization of the protein however, it is in the target DNA binding groove (Fig. S2D) and would block target DNA binding. After transposon DNA excision, the protein must capture a target DNA for integration into a new location. For that, the alpha helices must rearrange and move out of the target DNA binding groove to allow the transposase to continue to the forward transposition reaction. This could be more evidence that DNA transposition must involve significant protein rearrangement. DmTNP also has extensions at the C-terminus (17aa) but they are not resolved in the cryo-EM structure, as presumably they are disordered.

The data presented here sheds light on the activities of the THAP9 proteins, including the similarities and differences of THAP9 to *Drosophila* P element transposase and raise interesting questions regarding the ability of the proteins to act on different transposon DNA ends. Further biochemical and structural studies on THAP9 proteins from zebrafish and other homologs will address these questions and contribute to a mechanistic understanding of how these protein-DNA complexes function. These efforts will contribute to our understanding of the effects of transposable elements in genome evolution and gene expression programs.

## Acknowledgments

We are grateful to William Thomas for his assistance in generating the zfTHAP9 model using Alphafold2 (Jumper et al., 2021). We also thank members of the Rio lab for suggestions and advice throughout the course of this work. This work used the Vincent J. Coates Genomics Sequencing Laboratory at the University of California, Berkeley, supported by National Institutes of Health (NIH) S10 Instrumentation grants S10RR025622, S10RR029668, and S10RR027303. This work was supported by NIH R35GM118121.

## Author contributions

Nitzan Kutnowski carried out the molecular biology, *in-vivo* assays and biochemical experiments. George E. Ghanim performed Alphafold2 modeling. George Ghanim and Yeon Lee provided technical advice and assistance on cell culture and transposase *in-vivo* assays. All authors wrote the manuscript.

## Conflict of interest statement

**none declared.**

## MATERIALS & METHODS

### Structured Methods

#### Reagents and Tools Table

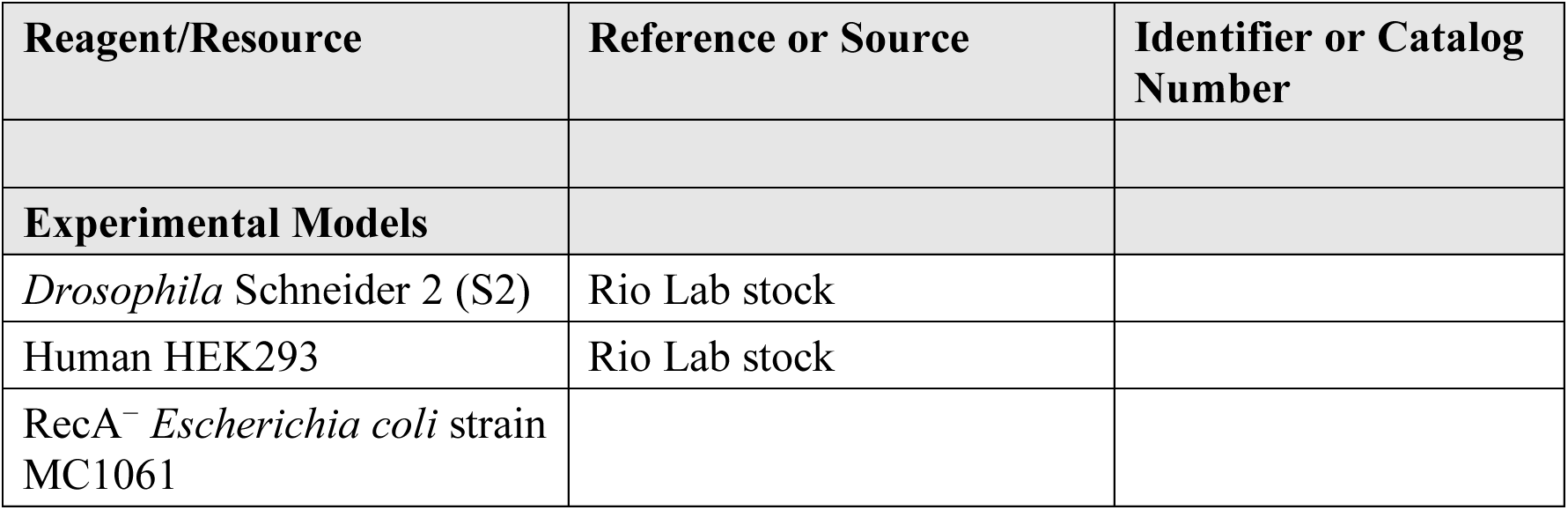
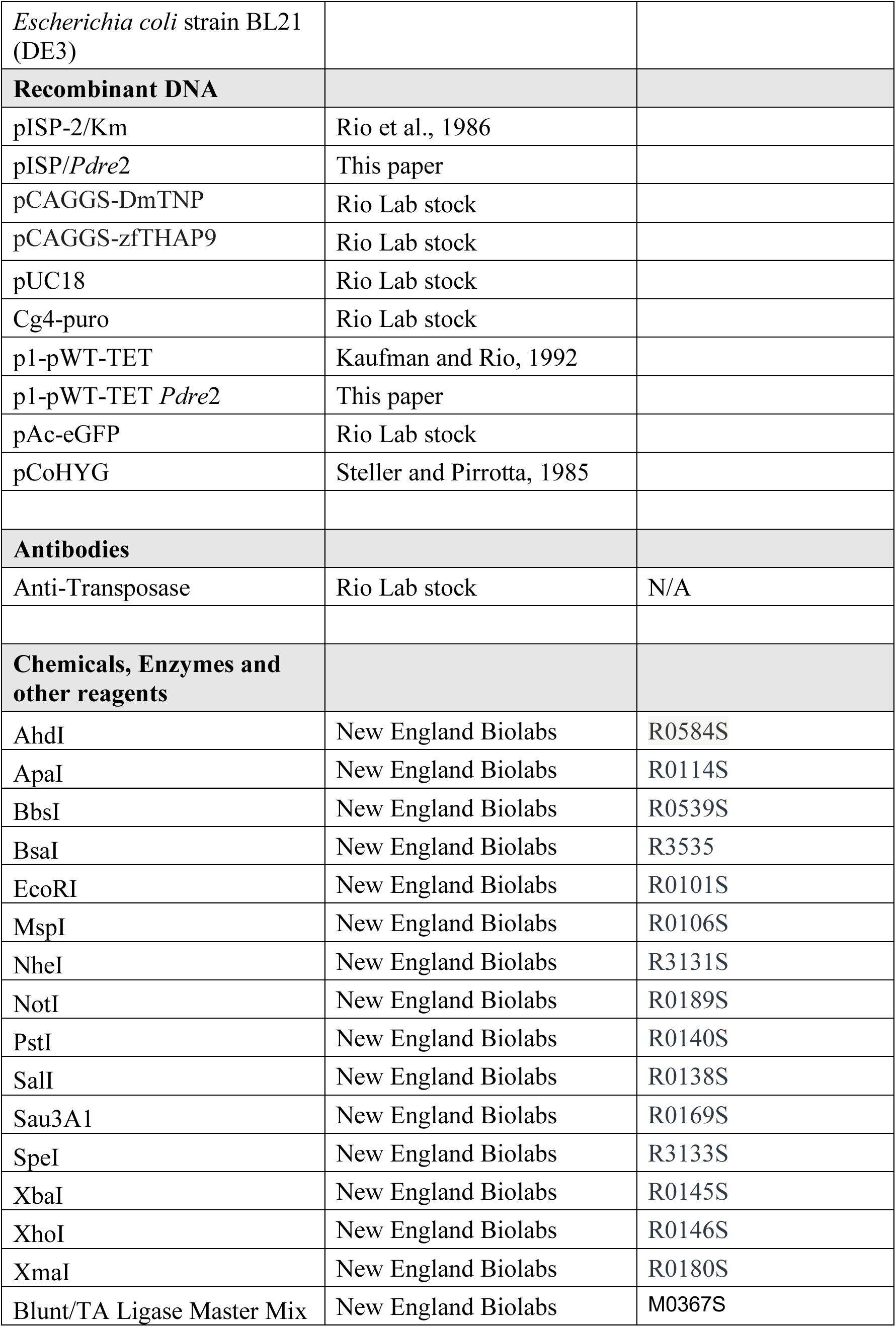
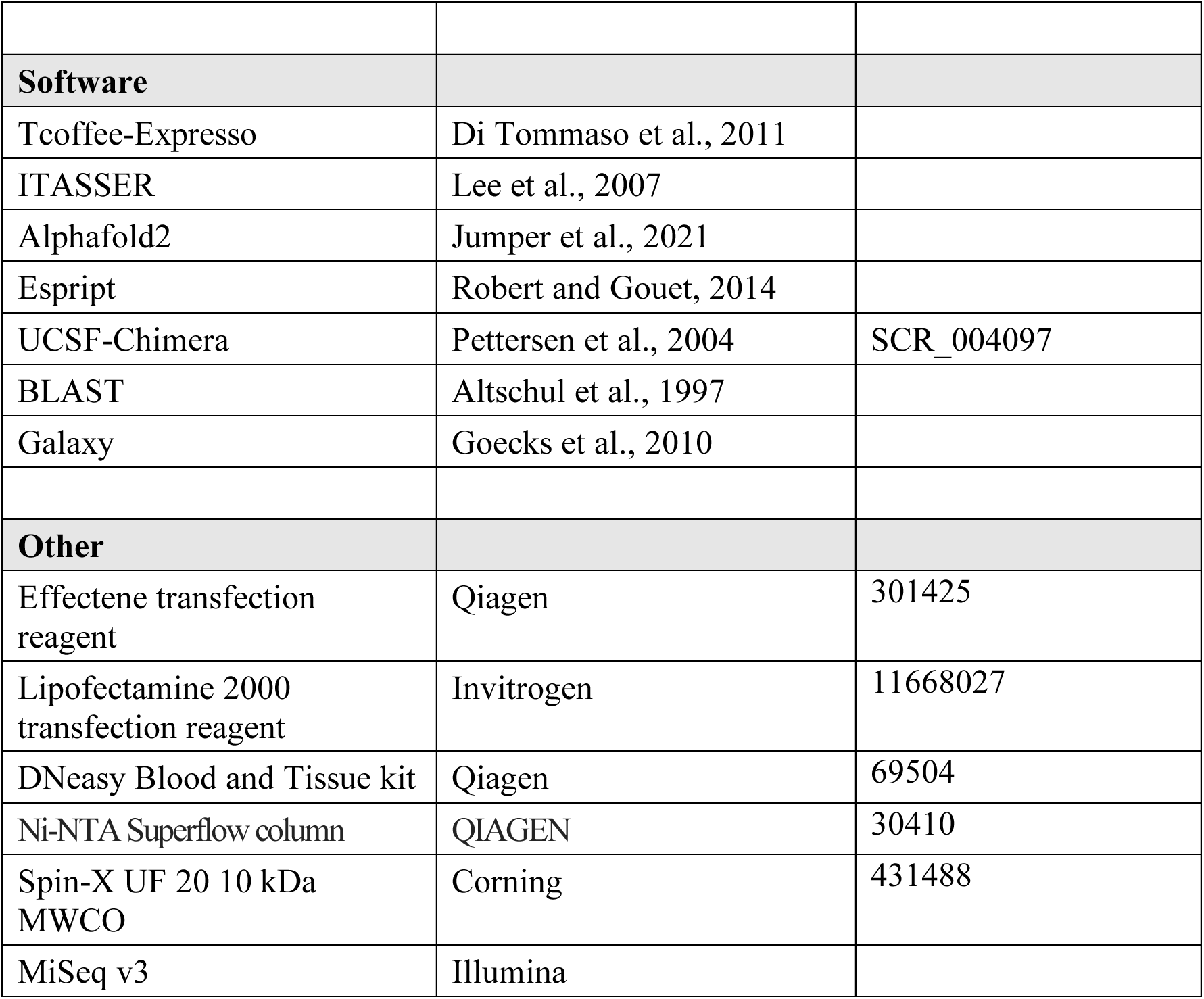

#### Bioinformatics analysis

All protein sequences were extracted from the NCBI website (http://www.ncbi.nlm.nih.gov/). Tcoffee-Expresso (Di Tommaso et al., 2011) was used to carry out structural sequence alignments of the DmTNP structure with the model of zfTHAP9 that generated using ITASSER (Lee et al., 2007) or Alphafold2 (Jumper et al., 2021). Multiple sequence alignments are displayed using Espript (Robert and Gouet, 2014). Molecular models were rendered using UCSF-Chimera (Pettersen et al., 2004). AlphaFold2 predictions were performed through a local installation of Colabfold 1.2.0 (Mirdita et al., 2022), running MMseqs2 (Mirdita et al., 2019) for homology searches and AlphaFold2-Multimer (Evans et al., 2021) for the prediction.

### Cell lines and culture

*Drosophila* Schneider 2 (S2) cells were obtained from long-term Rio Lab stock and grown in M3 media supplemented with 5% fetal bovine serum. Human HEK293 cells were obtained from long-term Rio Lab stock and grown in DME media supplemented with 10% fetal bovine serum. The cells were tested for mycoplasma in the MCB cell culture facility.

### *In vivo* excision assays

*In vivo* excision assays were performed using the pISP-2/Km (Rio et al., 1986) (kanamycin-sensitive because a 0.6 kb P element insertion disrupts the kanamycin resistance open reading frame), pISP/*Pdre*2 (kanamycin-sensitive because a 1.1 kb zebrafish *Pdre*2 insertion disrupts the kanamycin resistance open reading frame) reporter plasmid and DmTNP or zfTHAP9 plasmid DNA (cloned into pPAc for S2 cells or pCAGGS for HEK293 cells).

For *Drosophila* Schneider 2 cells, 1 × 10^6^ cells were transfected with 1µg pISP/2Km or pISP/*Pdre*2 reporter plasmid and either 0.25 µg empty plasmid (pUC18) or transposase source, using Effectene transfection reagent (Qiagen, Germantown, MD).

In HEK293 cells, 1 × 10^6^ cells were transfected with 1µg pISP-2/Km or pISP/*Pdre*2 reporter plasmid and either 1 µg empty plasmid (pUC18) or transposase source, using Invitrogen Lipofectamine 2000 transfection reagent (Invitrogen). At 24 hr after transfection, cells were washed with PBS, then harvested for immunoblot analysis and plasmid DNA recovery. Plasmid DNA was recovered as previously described (Mul and Rio, 1997), resuspended in 10 µl TE buffer (10 mM Tris-HCl pH 7.5, 1 mM EDTA), then 1 µl was used to transform RecA^−^ *Escherichia coli* strain MC1061 with a Bio-rad Gene Pulser as described by the manufacturer. Cells were grown for 2 hr at 37 °C with shaking, then plated onto Luria broth plates containing either 100 µg/ml of ampicillin or 100 µg/ml of ampicillin and 50 µg/ml of kanamycin. Colonies were allowed to grow for 16 hr at 37°C, then counted.

Excision activity was calculated by dividing the total number of kanamycin and ampicillin colonies by the total number of ampicillin colonies. “Relative integration activity” was calculated by computing activity as a percentage of the activity of the wild-type protein. DNA plasmid extracted from the ampicillin and kanamycin-resistant bacterial colonies was sequenced and digested with *PvuII* to verify excision.

### Integration assay in HEK293 cells

Integration assays were performed by transfecting a reporter plasmid, Cg4-puro (carries an SV40 promoter-puromycin gene inside the Cg4 P elements or zebrafish *Pdre*2 elements) and the DmTNP/zfTHAP9/pUC18 (negative control) plasmid DNA into 1 × 10^6^ HEK293 cells plated per well of a 6-well plate using Lipofectamine 2000 transfection reagent (Invitrogen). Each well was transfected with 2 µg of total plasmid DNA (50 ng Cg4-puro, +/- 0.5 µg DmTNP/zfTHAP9 expression plasmids and made up to 2 µg with an empty plasmid, (pUC18)). After 48 hours, the cells were trypsinized and re-plated on 10 cm dishes. Gene transfer events were monitored by selection with 2 µg/ml puromycin for two weeks. Puromycin-resistant colonies were stained with crystal violet and counted. “Relative integration activity” was calculated by computing activity as a percentage of the activity of DmTNP or zfTHAP9.

### Mapping P element integration sites using Splinkerette PCR

Genomic DNA was isolated from individual puromycin-resistant colonies obtained after integration, digested with restriction enzymes MspI or Sau3A1 and characterized by splinkerette PCR followed by DNA sequencing, as described previously (Potter and Luo, 2010).

### *In vivo* plasmid-to-plasmid transposition assay

For the *in-vivo* plasmid-to-plasmid transposition assay, S2 cells were co-transfected as above for excision experiments with 500 µg p1-pWT-TET (Kaufman and Rio, 1992)/ p1-pWT-TET *Pdre*2 (carries tetracycline resistance gene inside the zebrafish *Pdre*2 elements) donor plasmid, 500 µg target plasmid (pAc-eGFP) and either 0.25 µg empty plasmid (pUC18) or transposase source plasmids.

For *in-vivo* plasmid-to-plasmid transposition assay in HEK293 cells, plasmids were co-transfected as above for excision with 500 µg p1-pWT-TET/ p1-pWT-TET *Pdre*2 donor plasmid, 500 µg target plasmid (pCAGGS-eGFP) and either 1 µg empty plasmid (pUC18) or transposase source.

At 24 hr after transfection, cells were washed with PBS, then harvested for plasmid DNA recovery. Plasmid DNA was recovered as previously described (Mul and Rio, 1997), resuspended in 10 µl TE buffer (10 mM Tris-HCl pH 7.5, 1 mM EDTA), then 1 µl was used to transform RecA^−^ *Escherichia coli* strain MC1061 with a Bio-rad Gene Pulser as described by the manufacturer. Cells were grown for 2 hr at 37°C with shaking, then plated onto Luria broth plates containing either 100 µg/ml of ampicillin or 100 µg/ml of ampicillin and 50 µg/ml of tetracycline. Colonies were allowed to develop for 16 hr at 37°C, then counted.

DNA plasmid extracted from the ampicillin and tetracycline-resistant bacterial colonies was sequenced and digested with *Bsa*I to verify integration.

### Plasmid rescue assay

For plasmid rescue assays, *Drosophila* Schneider Line-2 cells were co-transfected as above for excision with 0.25 µg p6.1 in pCoHYG/ p6.1 *Pdre*2 in pCoHYG (Steller and Pirrotta, 1985), 250 µg transposase source and empty plasmid (pUC18). After one day of transfection, cells were split to T25 flask and selected with 200 µg/ml hygromycin for one week.

For plasmid rescue assay HEK293 cells were co-transfected as above for excision with 0.25 µg P6.1 in PGK PURO/ P6.1 in *Pdre*2 PGK PURO (Steller and Pirrotta, 1985) donor plasmid, 500 ng transposase source and empty plasmid (pBSK+ tetramer). After two days of transfection, cells were trypsinized and re-plated on 10 cm dish and selected with 2 µg/ml puromycin for two weeks.

Selected cells were harvested for genomic DNA preparation using DNeasy Blood and Tissue kit (Qiagen, Germantown, MD). 2 µg of genomic DNA was digested with one of few combinations of restriction enzymes (Table S5, all enzymes from NEB, Ipswich, MA). Digested genomic DNA was ligated using T4 ligase (NEB, Ipswich, MA) at 16°C for 16 hr. ligated DNA was precipitated and resuspended in 30 µl. One µl was used to transform RecA^−^ *Escherichia coli* strain MC1061 with a Bio-rad Gene Pulser as described by the manufacturer. Cells were grown for 1.5 hr at 37°C with shaking, then plated onto Luria broth plates containing 50 µg/ml of kanamycin. Colonies were allowed to develop for 16 hr at 37°C. Plasmid DNA was isolated from individual kanamycin-resistant colonies followed by DNA sequencing using primers that read through the TIR elements of the *Drosophila* P element transposon or zebrafish *Pdre*2 elements to verify transposition. The reads were aligned to the human or *Drosophila* genome using BLAST (Altschul et al., 1997).

### zfTHAP domain expression and purification

The gene encoding zfTHAP domain was cloned into expression vector to encode a version of the protein bearing a N-terminal MBP followed by a His6-tag. For protein expression, the plasmid was introduced into *Escherichia coli* strain BL21 (DE3). The cells were grown at 37°C and shaken at 175 rpm. IPTG (1 mM) was added at an OD_600_ of 0.6 and the cells were grown for 16 hr at 18°C. The cells were harvested by centrifugation (5000 x g) for 10 min. at 4°C. Harvested cell pellets were washed with PBS and snap-frozen in liquid nitrogen for later purification.

Cell pellets were thawed on ice, disrupted in 40 ml lysis buffer (25 mM HEPES-KOH, pH 7.6, 0.5 M KCl, 10 mM ZnCl_2_, 20 mM imidazole, 0.5 mM TCEP, 1 mM PMSF and 1X protease inhibitor cocktail), briefly sonicated, then clarified by centrifugation at 25,000 x g for 30 min. Supernatants were filtered through a 0.22 µm syringe filter before application to 5 ml of nickel resin (Ni-NTA Superflow, QIAGEN) equilibrated in loading buffer (25 mM HEPES-KOH pH 7.6, 0.5 M KCl, 10 mM ZnCl_2_, 20 mM imidazole, 0.5 mM TCEP and 1 mM PMSF). The resin was washed with 13 CVs loading buffer. Protein was eluted with a linear gradient of 20 mM to 0.5 M imidazole over 10 CVs. The eluted protein was diluted into a low-salt buffer (25 mM HEPES-KOH pH 7.6, 0.1 M KCl, 10 mM ZnCl_2_, 0.5 mM TCEP and 1 mM PMSF), then loaded onto a 1 ml HiTrap heparin HP column (GE Healthcare) pre-equilibrated in low salt buffer and eluted with a linear gradient of 100 mM to 1 M KCl over 25 CVs.

Peak fractions were concentrated to 1 mg/ml using a Spin-X UF 20 10 kDa MWCO (Corning). Protein concentration was determined by UV at wavelength of 280 nm. The final protein purity was determined by SDS–PAGE and Coomassie staining to be around 85% (Fig. S9).

### SELEX experiments

To identify DNA sequences recognized by zfTHAP domain by SELEX, a library of sequences was generated. The initial SELEX library was a pool of synthesized oligonucleotides comprising 20 bp-long randomized DNA fragments flanked by 20 bp-long primer sites for PCR amplification. This initial library was prepared using 14 PCR cycles. 20 µl purified zfTHAP domain (1 mg/ml) was bound to 50 ml Ni ^+^ NTA beads, equilibrated and washed three times in 200 µl binding buffer (25 mM HEPES-KOH, pH 7.6, 0.1 M KCl, 10 mM ZnCl_2_ and 0.5 mM TCEP), prior to the addition of DNA sequences (1 µg) from the initial library. The zfTHAP domain–DNA binding reaction was carried out at room temperature for 5 min. by rocking the beads in an Eppendorf tube. After washing three times with 200 µl binding buffer, the protein and bound DNA were eluted and separated using 200 µl elution buffer (25 mM HEPES-KOH, pH 7.6, 0.1 M KCl, 10 mM ZnCl_2_, 300 mM imidazole and 0.5 mM TCEP). Aliquots (2 µl) from the eluted fractions were subjected to 14 cycles of PCR amplification. Then, the DNA fragments served as input for another round of selection. Products of SELEX rounds were sequenced on a MiSeq v3 machine (Illumina) with 150-bp single reads sequencing at the Vincent J. Coates Genomics Sequencing Laboratory at the University of California, Berkeley. Bioinformatics and statistical analyses, as well as visualization of the results, were carried out using the Galaxy package and server (Goecks et al., 2010).

## Notes

### Competing Interest Statement

The authors have declared no competing interest.

