## Supplementary data for "Activity of zebrafish THAP9 transposase and zebrafish P element-like transposons"

### Figure S1

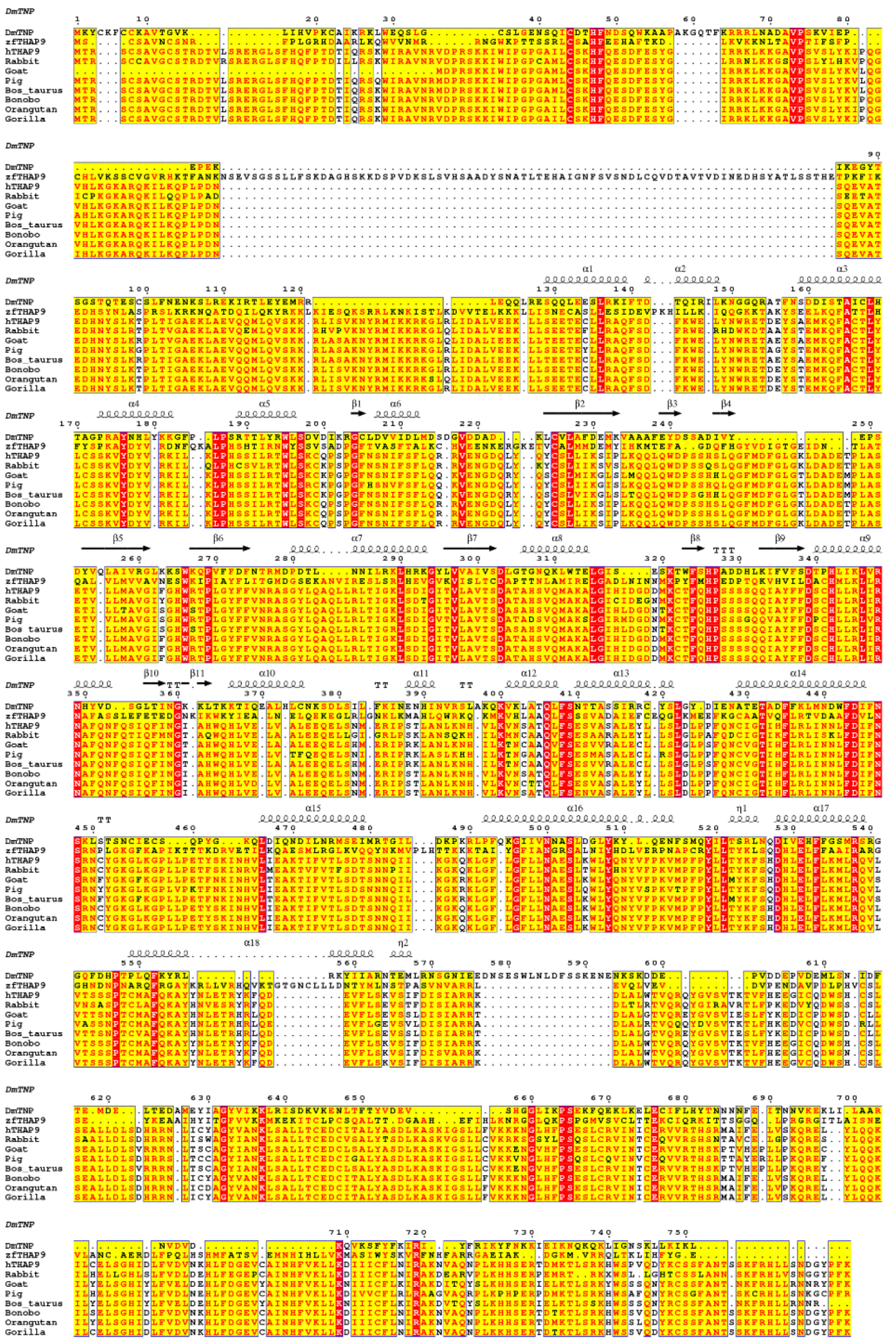

**Fig. S1: Multiple sequence alignment (MSA) of the *Drosophila* P element transposase (DmTNP) with various THAP9 proteins.**

Secondary structure elements are shown according to the DmTNP structure. THAP9 proteins: *Drosophila* P element transposase (DmTNP), zebrafish P element transposase (zfTHAP9), human P element transposase (hTHAP9) with homologous proteins from rabbit, gallus, goat, pig, Bos taurus, bonobo, orangutan and gorilla. Amino acids in red boxes are identical, while those in yellow boxes are conservative substitutions. The sequences were aligned using Tcoffee (Di Tommaso et al., 2011) and displayed to show secondary structure elements, as determined by ESPript3.0 (Robert and Gouet, 2014).

**Figure S2**

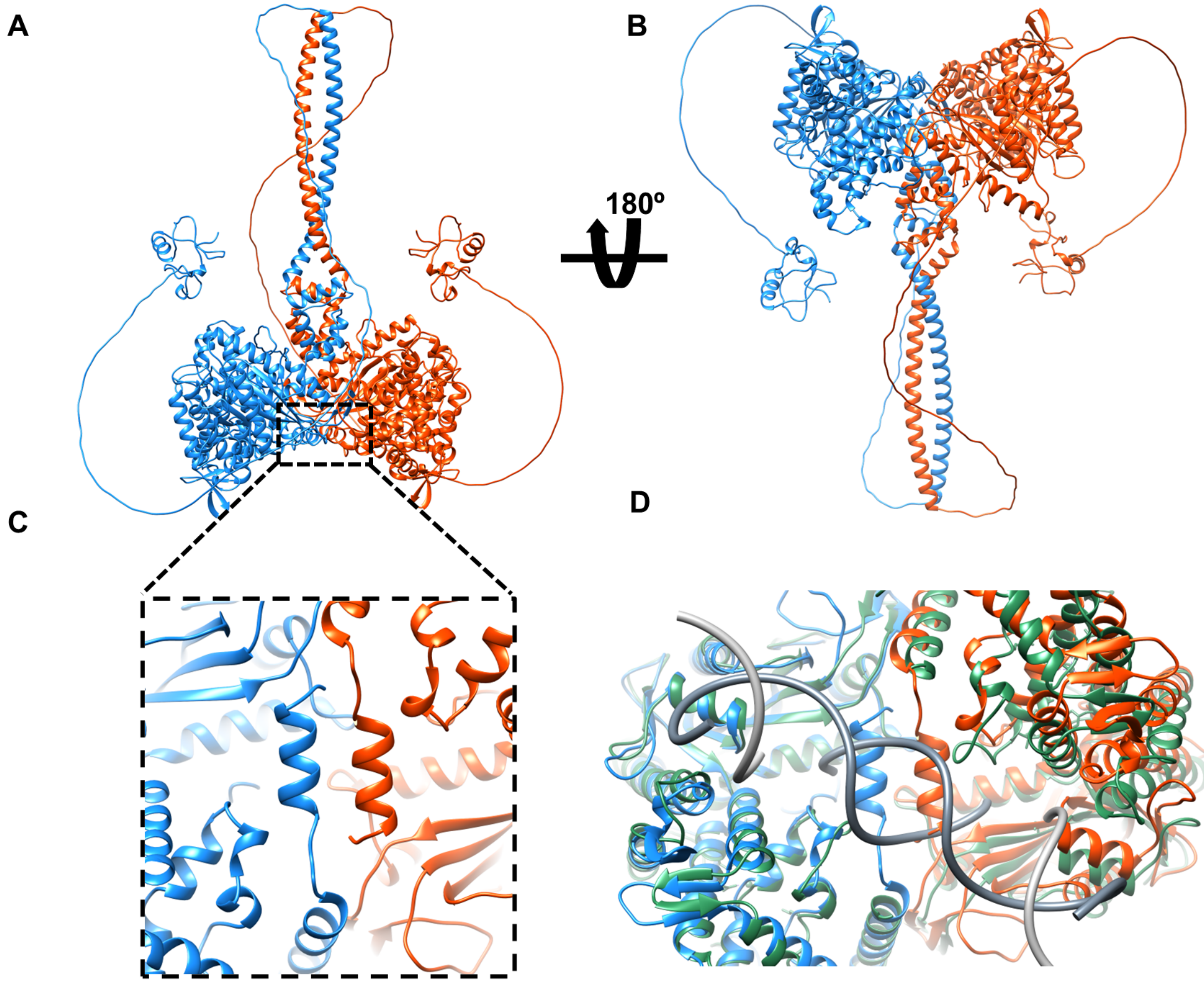


**Figure S2. AlphaFold2 structural prediction of the zfTHAP9 homodimer.**

**A and B.** Two different views of an AlphaFold2 structural prediction of the zfTHAP9 homodimer. Each subunit is colored blue or orange. **C.** Close-up view of the pair of alpha helices close to the end of zfTHAP9 (R920-F929). **D.** Close-up view of the superposition of the Drosophila P element transposase (DmTNP) cryo-EM structure (green) (Ghanim et al., 2019) and the zfTHAP9 model (blue and orange). The target DNA is colored in gray.

**Figure S3**

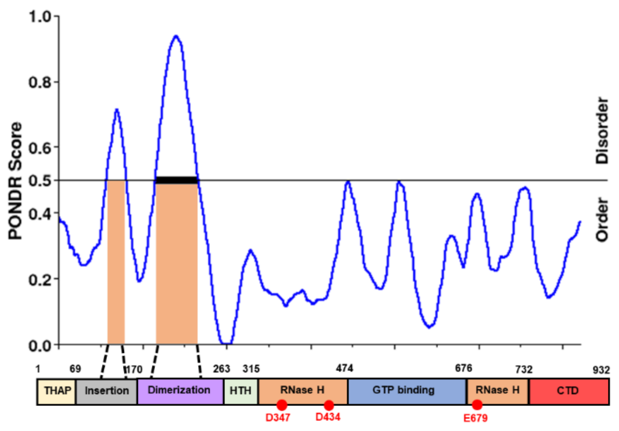


**Fig. S3: Prediction of disordered regions in the zebrafish THAP9 protein.**

PONDR scores of predicted disordered regions. Disordered regions that were predicted with high confidence (indicated by black bars), are within the leucine zipper dimerization domain and the insertion domain of zfTHAP9.

**Figure S4**

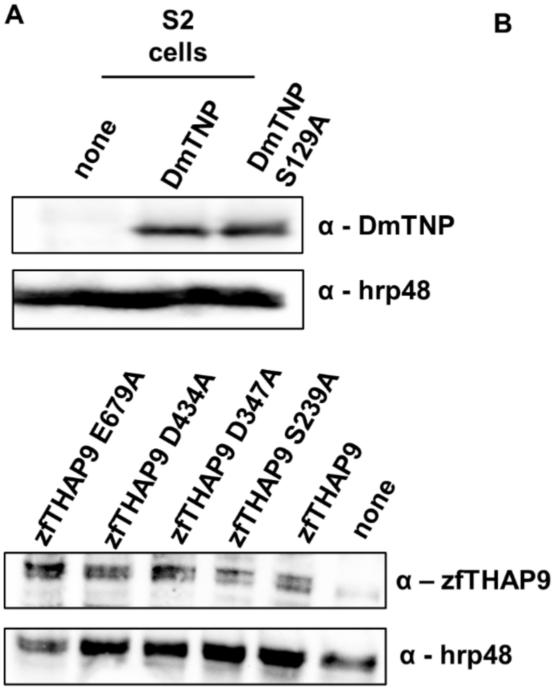


**Fig. S4: Immunoblots of DmTNP and zfTHAP9 proteins expressed after transfection of** ***Drosophila*** **S2 and human HEK293 cells.**

**A.** Representative immunoblots of wild type and mutant transposase protein expression levels in S2 cells. Cells were harvested 24hr after transfection. Membrane was cut and then immunoblotted with anti-transposase antibodies (α-TNP) or a loading control (α-HRP48).

**B.** Representative immunoblots of transposase protein expression levels in HEK293 cells. Cells were harvested 24hr after transfection.

**Figure S5**

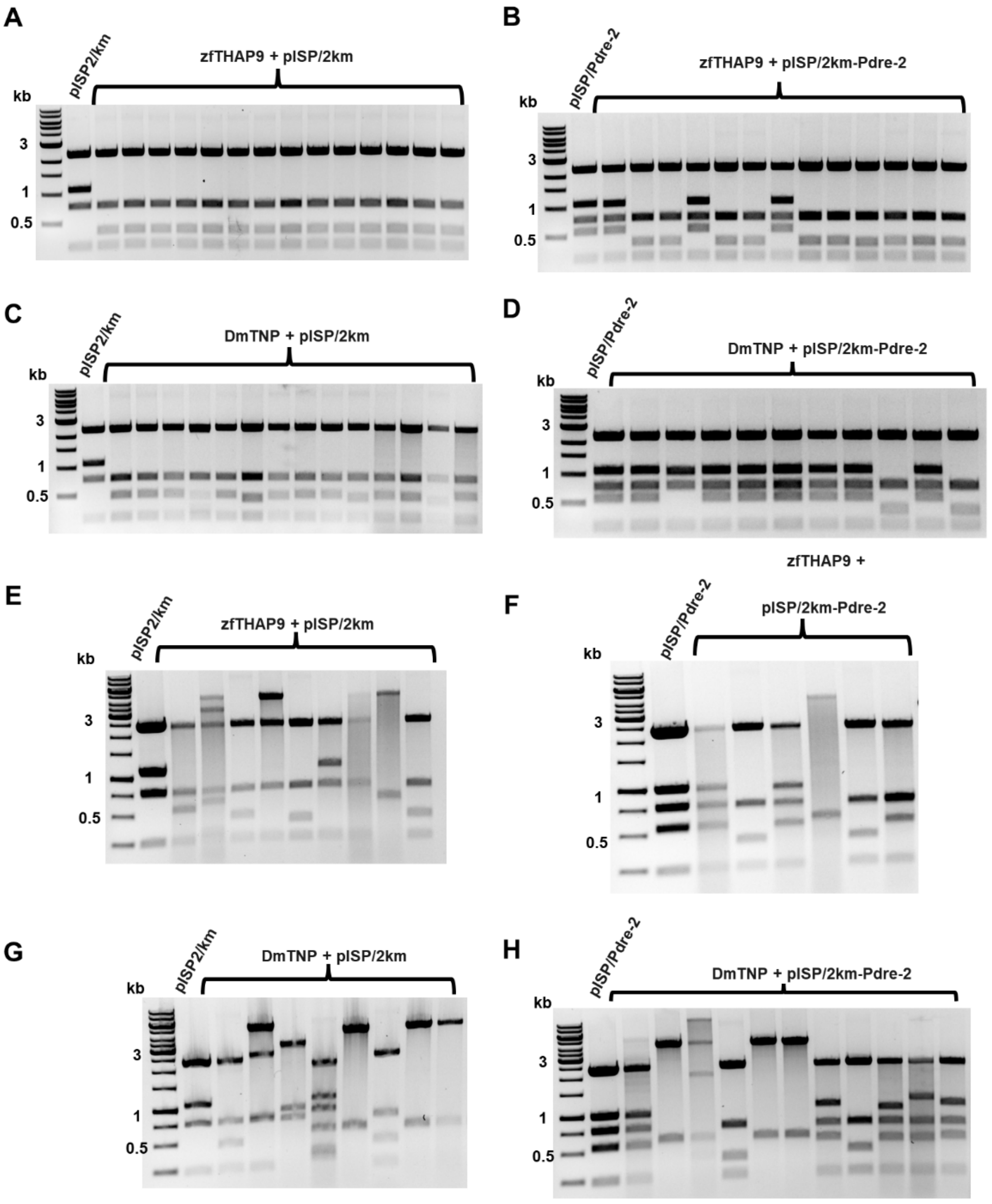


**Fig. S5: Analysis of excised plasmid DNA derived from *Drosophila* P element or *Pdre*2 element excision events recovered from *Drosophila* S2 cells and human HEK293 cells.**

Photographs of ethidium bromide-stained 1% agarose gels showing the *Pvu*II cleavage patterns of the pISP/2Km or pISP/*Pdre*2 derived plasmid DNAs that were recovered from single kanamycin and ampicillin-resistant bacterial colonies obtained after transfection of *Drosophila* S2 cells (**A-D**) or HEK293 cells (**E-H**) with the DmTNP or zfTHAP9 expression vectors and the indicated reporter plasmids. *Pvu*II restriction endonuclease cleavage of the unmodified pISP/2Km reporter plasmid gives rise to fragments of 2 kb, 1.1 kb, 0.75kb and 0.25kb fragments, whereas excision of the P element generates an approximately 0.5 kb *Pvu*II cleavage product, instead of the 1.1kb fragment, depending on the deletions that occur flanking the break site. *Pvu*II restriction endonuclease cleavage of the unmodified pISP/*Pdre*2 reporter plasmid gives rise to five fragments of 2.7 kb, 0.75kb, 1.1kb, 0.5kb, and 0.25kb, whereas excision of the *Pdre*2 element generates only four fragments so the 1.1kb fragment is eliminated, depending on the deletions that occur flanking the break site.

### Figure S6

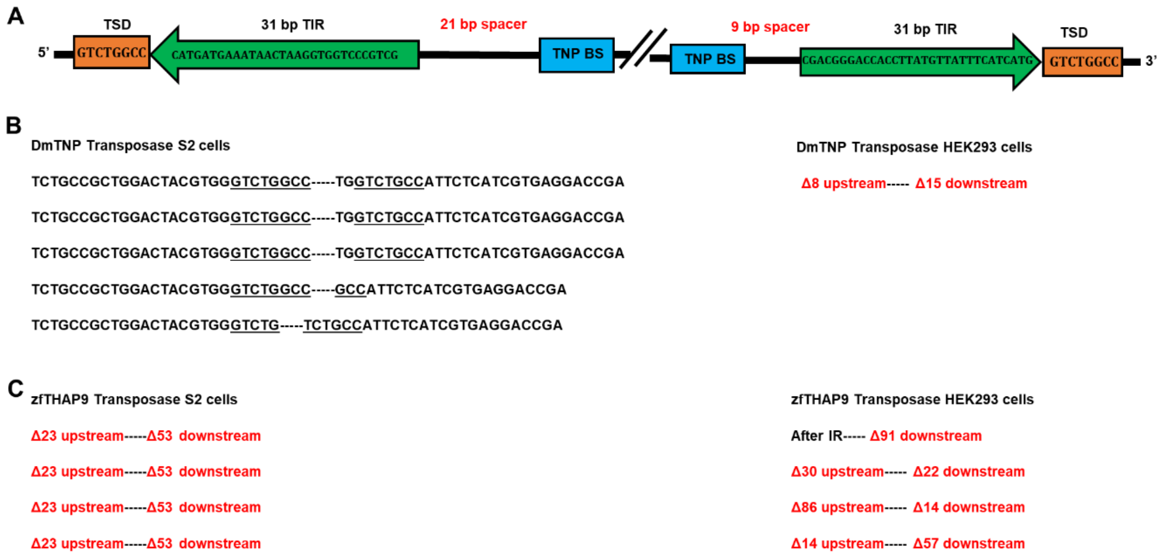


**Fig. S6: DNA sequence analysis of pISP/2Km excision products from *Drosophila* S2 and human HEK293 cells.**

**A.** Organization of the *Drosophila* P elements. Indicated are the 8 bp target site duplication (TSD), 31bp terminal inverted repeat (TIR) and TNP binding sites. **B-C.** Left: Examples of P elements donor site repair sequence junctions that were found in plasmid DNA prepared from individual kanamycin-resistant bacterial colonies obtained after electroporation of bacteria with low molecular weight DNA isolated from S2 cells. The 8 bp target site duplication (TSD) flanks the P elements insertion are underlined. Right: Examples of P element donor site repair junctions that were sequenced in plasmid DNA prepared from individual kanamycin-resistant bacterial colonies obtained after electroporation of bacteria with low molecular weight DNA isolated from HEK293 cells.

### Figure S7

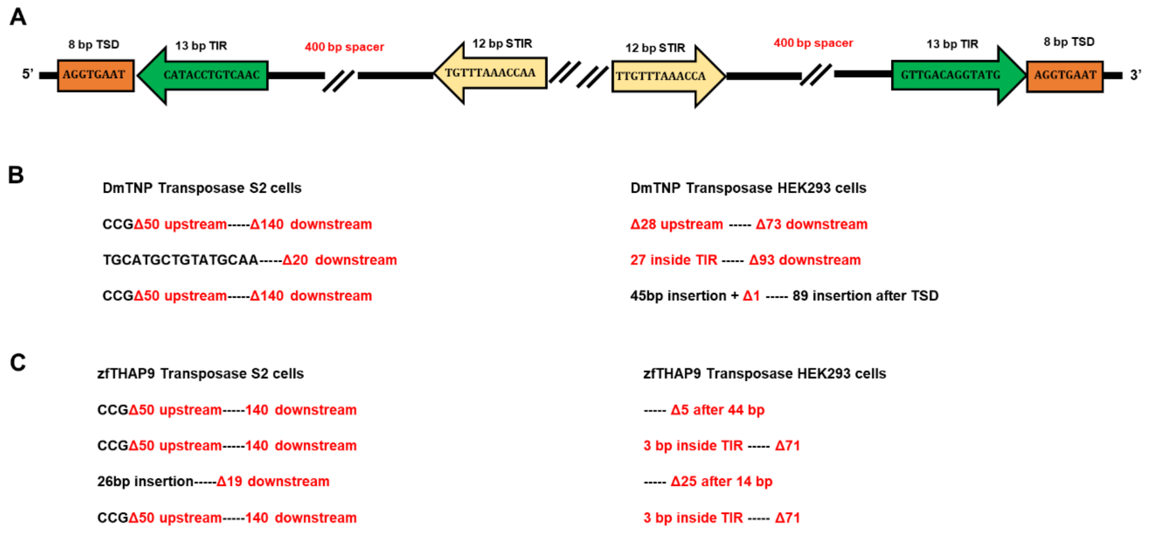


**Fig. S7: DNA sequence analysis of pISP/*Pdre*2 excision products from *Drosophila* S2 and human HEK293 cells.**

**A.** Organization of the zebrafish *Pdre*2 elements. Indicated are the 8 bp target site duplications (TSD), 13 bp terminal inverted repeats (TIR) and 12 bp internal inverted repeats. **B-C.** Left: Examples of *Pdre*2 donor site repair junctions that were sequenced in plasmid DNA prepared from individual kanamycin-resistant bacterial colonies obtained after electroporation of bacteria with low molecular weight DNA isolated from S2 cells. Right: Examples of *Pdre*2 donor site repair junctions that were sequenced in plasmid DNA prepared from individual kanamycin-resistant bacterial colonies obtained after electroporation of bacteria with low molecular weight DNA isolated from HEK293 cells.

### Figure S8

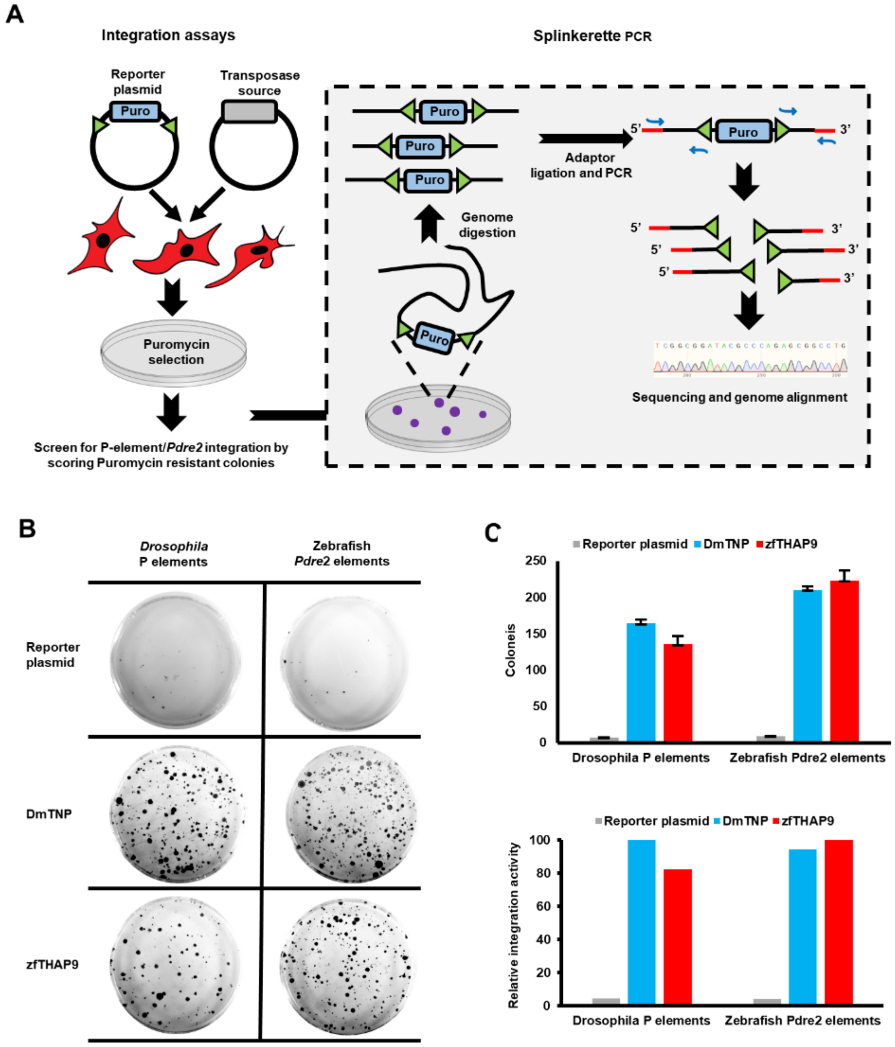


**Fig. S8: Integration assays followed by splinkerette PCR (spPCR) in HEK293 cells.**

**A.** Schematic diagram of P elements integration assays followed by spPCR.

**B.** Crystal-violet staining of colonies obtained after two weeks puromycin selection of HEK293 cells co-transfected with the reporter plasmid and DmTNP, zfTHAP9 expression vectors or pUC18 (negative control). **C.** Upper panel: A graph showing the average number of colonies of three independent experiments. Grey – negative control, blue – DmTNP and red – zfTHAP9. Lower panel: A graph comparing P elements or *Pdre*2 elements transposition activity. The relative activity was calculated compared to DmTNP or zfTHAP9.

**Figure S9**

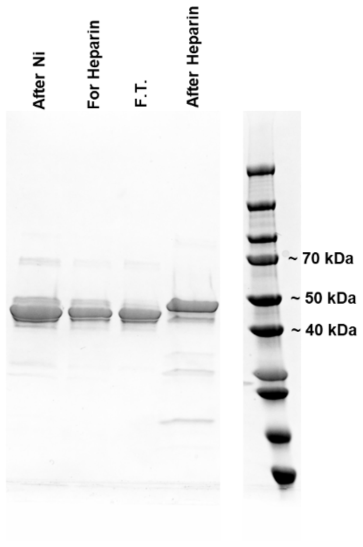


**Fig. S9: Purification of zfTHAP DNA-binding domain.**

SDS-PAGE of the purification process of MBP-zfTHAP DNA-binding domain. First lane: sample after $Ni^{2+}$ column, second lane: sample for heparin column, third lane: flow-through, forth lane: heparin column elution.

**Table S1:** Catalytic residues

| DmTNP | zfTHAP9 |
|---|---|
| D230 | D347 |
| D347 | D434 |
| E531 | E679 |

**Table S2:** GTP binding residues

| | | | | | | | | | |
|---|---|---|---|---|---|---|---|---|---|
| **DmTNP** | K385 | K400 | V401 | S409 | N440 | F443 | D444 | N447 | D528 |
| **zfTHAP9** | K522 | K536 | V537 | S545 | N578 | F581 | D582 | N585 | D676 |

**Table S3:** Excision assays results in S2 cells

| Plasmids transfected | Excision activity (×10$^6$) |
|---|---|
| **pISP/2Km** | |
| **Negative control** | 0.25±0.06 |
| **DmTNP** | 7.66±0.32 |
| **DmTNPS129A** | 10.76±0.32 |
| **zfTHAP9** | 2±0.55 |
| **zfTHAP9S239A** | 2.97±0.18 |
| **zfTHAP9D347A** | 0.95±0.24 |
| **zfTHAP9D434A** | 0.72±0.11 |
| **zfTHAP9E679A** | 0.33±0.11 |
| **pISP/*Pdre*2** | |
| **Negative control** | 0.12±0.08 |
| **DmTNP** | 2.54±0.48 |
| **DmTNPS129A** | 5.11±1.49 |
| **zfTHAP9** | 2.9±0.13 |

**Table S4:** Plasmid-to-Plasmid results

| Plasmids transfected | Integration activity ($\times 10^6$) |
|---|---|
| **Negative control** | 4.01±1.93 |
| **DmTNP** | 149.19±15.13 |
| **DmTNPS129A** | 188.41±12.8 |

**Table S5:** restriction enzymes used for plasmid rescue experiments:

| | |
|---|---|
| ***Drosophila* P elements- S2 cells *Ahd*I + *Apa*I** | |
| 1 | *Spe*I + *Nhe*I |
| 2 | *Xba*I + *Nhe*I |
| 3 | *Sal*I + *Nhe*I + *Xho*I |
| 4 | *Eco*RI + *Bbs*I |
| 5 | *Bsa*I + *Bbs*I+ *Spe*I+ *Nhe*I |
| ***Drosophila* P elements- HEK293 cells *Ahd*I + *Apa*I + *Bbs*I** | |
| 1 | *Xba*I + *Nhe*I |
| 2 | *Pst*I |
| 3 | *Xba*I + *Sal*I |
| **Zebrafish *Pdre*2 S2- cells *Ahd*I + *Bsa*I+ *Bbs*I** | |
| 1 | *Xba*I + *Spe*I |
| 2 | *Xma*I |
| 3 | *Spe*I + *Nhe*I |
| **Zebrafish *Pdre*2- HEK293 cells *Ahd*I + *Bsa*I** | |
| 1 | *Bsa*I |
| 2 | *Not*I |
